# Spike-specific IgG4 generated post BNT162b2 mRNA vaccination is inhibitory when directly competing with functional IgG subclasses

**DOI:** 10.1101/2025.02.12.637095

**Authors:** Jerry C. H. Tam, Abbie C. Sibayan, Jeffrey Seow, Carl Graham, Ashwini Kurshan, Blair Merrick, Richard J. Stanton, Katie J. Doores

**Affiliations:** Department of Infectious Diseases, School of Immunology & Microbial Sciences, King’s College London, London, UK; Centre for Clinical Infection and Diagnostics Research, Department of Infectious Diseases, Guy’s and St Thomas’ NHS Foundation Trust, London, UK; Division of Infection and Immunity, School of Medicine, Cardiff University, Cardiff, United Kingdom

## Abstract

The rapid development of vaccines against SARS-CoV-2 during the COVID-19 pandemic proved vital in controlling viral spread and reducing mortality and morbidity. Both neutralising activity and effector function activity of Spike-specific antibodies have been shown to be important for their protective and therapeutic activity. However, several recent studies have reported that vaccination with mRNA based COVID-19 vaccines can lead to elevated levels of Spike-specific IgG4, an isotype which is often considered anti-inflammatory due to its reduced binding to Fcγ receptors on immune cells. Here we show that the level of Spike-specific IgG4 produced following BNT162b2 vaccination is impacted by the interval between and frequency of vaccines boosts, prior SARS-CoV-2 infection (hybrid immunity), breakthrough infection and bivalent vaccine boosters. Despite the increase in Spike-specific IgG4 between the 2^nd^ and 3^rd^ BNT162b2 vaccine dose, neutralisation, ADCD and ADCP activity all increased. Through expression of SARS-CoV-2 monoclonal antibodies cloned as IgG1, IgG2, IgG3 and IgG4, we demonstrated that whilst Spike-specific IgG4 had reduced effector function activity, including ADCC, ADCD and ADCP, IgG4 was only inhibitory when directly competing with functional IgG subclasses binding to an overlapping epitope. In the context of polyclonal plasma, ADCC and ADCD activity could not be depleted by addition of high concentrations of a Spike-specific IgG4 mAb cocktail suggesting the non-stimulatory effect of Spike-specific IgG4 may be hidden in more complex scenarios, such as polyclonal mixes in serum.

## Introduction

The COVID-19 pandemic led to the rapid development and deployment of several COVID-19 vaccines utilising a range of different vaccine platforms (Bok et al., 2021). Vaccines targeting SARS-CoV-2 Wuhan Spike glycoprotein have been highly effective in reducing morbidity and mortality from COVID-19 (Higdon et al., 2022). In particular, the pandemic led to the first widespread clinical use of mRNA based vaccines; including BNT162b2 from BioNTech/Pfizer (Polack et al., 2020) and Spikevax from Moderna (Baden et al., 2021). These were initially deployed in a 2-dose regimen, separated by a 3-week interval (Baden et al., 2021; Polack et al., 2020). However, the United Kingdom quickly changed to a 12-week interval between doses, with the rationale of ensuring some degree of SARS-CoV-2 immunity for a wider proportion of the population (UKHSA, 2022). To overcome waning antibody levels (Seow et al., 2020), a third dose was offered as a booster, which had the additional benefit of broadening the antibody response towards SARS-CoV-2 variants of concern (Graham et al., 2022; Kim et al., 2022; Roltgen et al., 2022; Sokal et al., 2021), as well as leading to an increased antibody binding avidity (Graham et al., 2022). More recently, updated mRNA vaccines based upon newer variants such as BA.1 and XBB.1.5 have been used as boosters (Kirsebom et al., 2023), not only for individuals primed with COVID-19 Wuhan-1 mRNA vaccines (Kirsebom et al., 2023; Lee et al., 2023), but also for individuals who had initially received other vaccines types, such as the AstraZeneca Chimpanzee Adenovirus Vector (ChAdOx) COVID-19 vaccine, AZD1222 (Akhtar et al., 2023). Given the safety and efficacy of COVID-19 mRNA vaccines, there has been increased interest in using this vaccine platform for more applications, including vaccines against other pathogenic viruses as well as against bacteria, and as immunotherapies for cancer treatment (Parhiz et al., 2024).

Whilst antibody binding and neutralisation correlate with protection against SARS-CoV-2 infection and severe COVID-19 disease (Goldblatt et al., 2022), the Fc region can also collaborate to enhance the protective capacity of antibodies. Animal studies have shown the importance of Fc-mediated effector functions of monoclonal antibodies (mAbs) for limiting SARS-CoV-2 immunopathology and viral persistence in animal models (Chan et al., 2021; Pierre et al., 2024; Schafer et al., 2021; Yamin et al., 2021), as well as robust antibody Fc-effector functions induced by vaccines correlating with protection in animal challenge studies (Gorman et al., 2021; Mercado et al., 2020; Tostanoski et al., 2020).

It has previously been reported that the use of COVID-19 mRNA-based vaccines can lead to skewing of the IgG subclass usage against SARS-CoV-2 Spike (Akhtar et al., 2023; Buhre et al., 2022; Gelderloos et al., 2024; Irrgang et al., 2023). In addition to the majority IgG1 and IgG3 response, Spike-specific IgG4 could also be detected at low levels in some individuals around 6-7 months post a second dose of either BNT162b2 or Spikevax (Akhtar et al., 2023; Irrgang et al., 2023) and at higher levels in the majority of individuals following a third dose of BNT162b2, as well as following a BNT162b2 booster vaccine in individuals who initially received the AZD1222 vaccine (Akhtar et al., 2023). This increase in Spike-specific IgG4 is notable, given that IgG4 is usually thought to be anti-inflammatory (Pillai, 2023), and is not typically observed following vaccination or infection (Urban et al., 1994). Instead, antigen-specific IgG4 has been reported in the context of allergy (Qin et al., 2022), autoimmunity (Motta and Culver, 2024; Rispens and Huijbers, 2023) and chronic inflammatory conditions (van de Veen et al., 2020). The perceived role of IgG4 as anti-inflammatory comes from its reduced ability to stimulate Fc-mediated antibody effector functions (Bruhns et al., 2009; Pillai, 2023), including antibody-dependent cellular cytotoxicity (ADCC), antibody-dependent complement deposition (ADCD) and antibody-dependent cellular phagocytosis (ADCP), due to the reduced affinity of the IgG4 Fc domain for FcγR receptors, in particular FcγRIIA, FcγRIIIA and FcγRIIIB (Vidarsson et al., 2014). IgG4 is the only immunoglobulin that can undergo Fab-arm exchange, a process in which half-molecules of IgG4 combine randomly resulting in bispecific immunoglobulins with reduced binding and/or neutralising activity due to a lower valency of binding (van der Neut Kolfschoten et al., 2007).

Antibody effector functions are important in clearing infection through binding of antibodies to antigens expressed on the surface of infected cells or to free virus (Beaudoin-Bussieres and Finzi, 2024). Effector functions have been shown to be important in both prophylactic and therapeutic protection models (Gorman et al., 2021; Zhang et al., 2023). For example, introduction of the LALA-PG mutations (L234A+L235A+P329G), which are known to reduce the binding of IgGs to FcγRs reduces the therapeutic activity of neutralising antibodies against SARS-CoV-2 (Winkler et al., 2021). Furthermore, non-neutralising Spike-specific IgG1 can confer protection in small animal challenge models if passively administered prior to challenge (Clark et al., 2024). Whilst the relevance of IgG subclass has not been studied *in vivo* in the context of SARS-CoV-2 infection in humans, antibody passive transfer experiments in the context of Influenza virus infection have revealed that IgG subclass is important in determining protection (DiLillo et al., 2014; Van den Hoecke et al., 2017). However, animal studies have limited utility for understanding the impact of increased levels of antigen-specific IgG4 given that it is only found in humans and great apes (Olivieri and Gambon Deza, 2018).

Whilst several studies report elevated levels of Spike-specific IgG4 following mRNA vaccination (Akhtar et al., 2023; Buhre et al., 2022; Gelderloos et al., 2024; Irrgang et al., 2023), there is an incomplete knowledge of factors that might influence the switch to IgG4 and how the level of Spike-specific IgG4 might be impacted by further mRNA vaccine booster doses and/or breakthrough infection (BTI). Furthermore, there is an incomplete understanding of the functional implications of the presence of Spike-specific IgG4 on the functional activity of the Spike-specific Total IgG. Here, we investigated the impact of; i) mRNA vaccine interval and frequency, ii) hybrid immunity, iii) breakthrough infection and iv) bivalent vaccines on the amount of IgG4 produced, the IgG4 binding specificity, breadth and avidity, and effector function activity in immune plasma. We observed that IgG4 production was impacted by the number of mRNA doses, the interval between these doses and SARS-CoV-2 infection (both pre- and post-vaccination). Through generation of a panel of class-switched Spike-specific mAbs, including IgG1, IgG2, IgG3 and IgG4, we demonstrate that whilst IgG4 poorly activates ADCC, ADCP and ADCD, it is capable of potent and equivalent neutralisation as IgG1. Importantly, IgG4 is only anti-inflammatory when competing directly for binding with an activating IgG1 but is functionally silent if non-competing. These findings provide important insights into the functional implications of elevated levels of Spike-specific IgG4 following BNT162b2 mRNA vaccination.

## Results

### Spike-specific IgG4 class-switching increases with multiple BNT162b2 vaccine exposures and time

To gain further understanding of the factors influencing class-switching to Spike-specific IgG4 during mRNA vaccination, we studied selected plasma from several UK based vaccine cohorts (**Table S1A** and **Fig. S1A**). These groups differed in; i) the spacing between the first and second vaccine doses, ii) prior or subsequent SARS-CoV-2 infection, and iii) use of a bivalent booster vaccine based on Wuhan-1/BA.1. We first measured the concentration of SARS-CoV-2 Spike-specific IgG in a subclass specific manner for each group using a semiquantitative ELISA (**Fig. 1**).

**Fig. 1.**
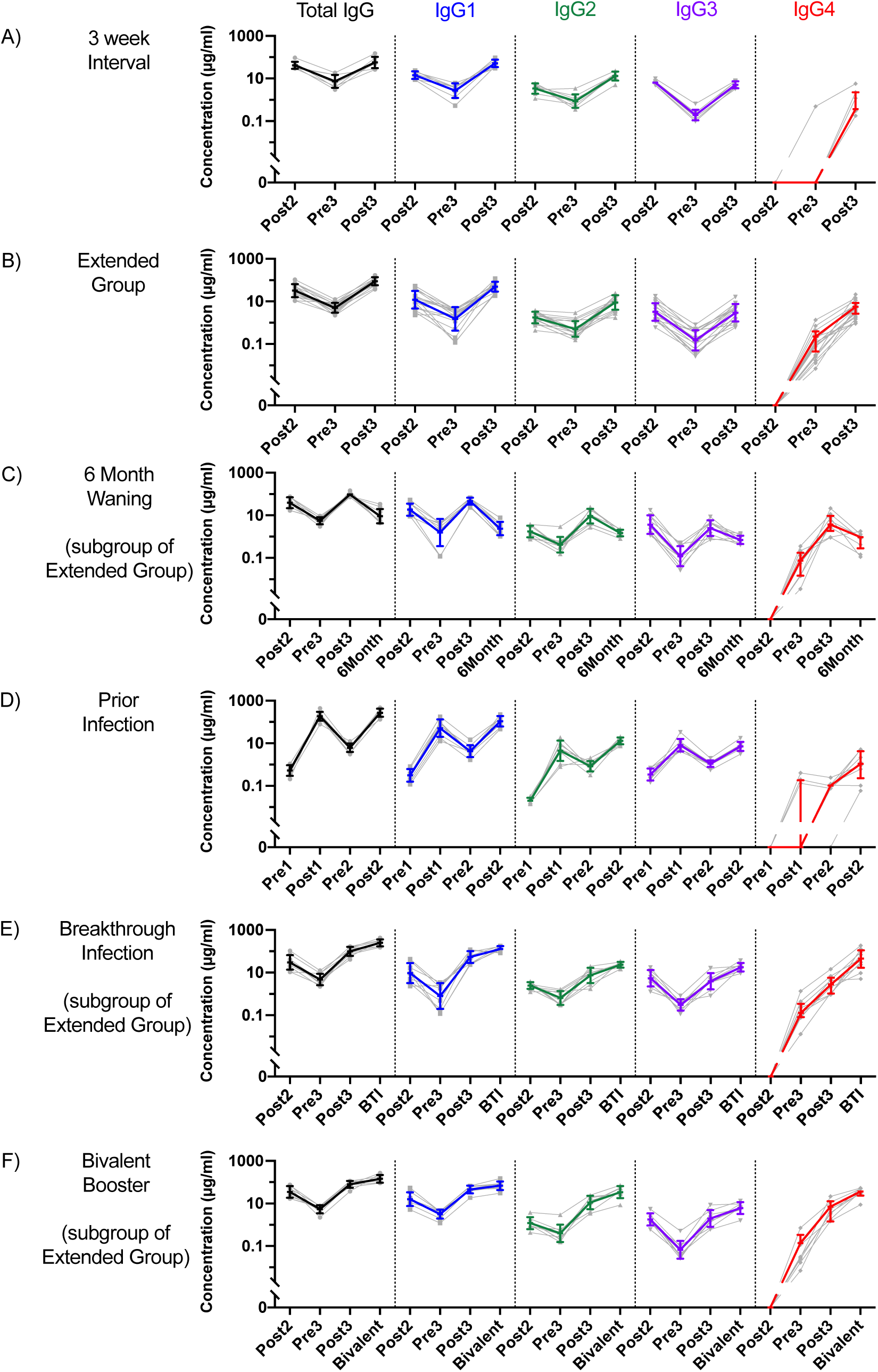
Spike-specific IgG4 production in individuals with different exposure histories. Semiquantitative ELISA measurement of Wildtype (WT) SARS-CoV-2 Spike-specific Total IgG (black), IgG1 (blue), IgG2 (green), IgG3 (purple) and IgG4 (red) for **(A)** 3 week interval between 1^st^ and 2^nd^ dose group (short group, n=8), **(B)** 12-week interval between 1^st^ and 2^nd^ dose group (extended group, n=17), **(C)** subgroup of extended group with sample a 6 months post third dose (n=9), **(D)** prior infection group (n=8), **(E)** subgroup of extended group with a breakthrough infection after three doses (n=9), **(F)** subgroup of extended group with a bivalent WT/BA.1 booster 1 year after third dose (n=8). Samples paired from an individual in grey with coloured data showing geometric mean and standard deviation. Pre1 = before 1^st^ dose, Post1 = 2 weeks after 1^st^ dose, Pre2 = before 2^nd^ dose, Post2 = 2 weeks after 2^nd^ dose, Pre3 = before 3^rd^ dose, Post3 = 2 weeks after 3^rd^ dose, 6Month = 6 month after 3^rd^ dose, BTI = 2 weeks after breakthrough infection, Bivalent = 2 weeks after bivalent dose. See Supplementary Table 1 for demographics and Supplementary Fig. S1 for precise timings. Semiquantitative ELISA limit of detection 0.05 µg/ml.

Following the standard BNT162b2 vaccination protocol (i.e. a 3–4-week interval between first and second dose, referred here to as “short group”), total IgG levels averaged 45.1 µg/ml after two mRNA doses (**Fig. 1A**). As previously reported, the total Spike-specific IgG levels waned over time, such that at six months post vaccination, they decreased to 9.05 µg/ml. A third vaccine dose was able to increase the level back to 67.0 µg/mL. This pattern of IgG levels can largely be explained by changes in IgG1, IgG2 and IgG3, which all follow a similar pattern. However, Spike-specific IgG4 only became detectable after the third dose of vaccine, similar to that previously reported (Gelderloos et al., 2024; Irrgang et al., 2023) with levels reaching 1.47 µg/ml.

In the UK, the majority of individuals receiving the BNT162b2 vaccine did so with an extended 8–12-week interval between the first and second doses (referred to here as “extended group”). The trends in total IgG, IgG1, IgG2 and IgG3 levels were similar to that of the short group (**Fig. 1B**). However, in contrast to the short group, Spike-specific IgG4 could be observed earlier, with Spike-specific IgG4 being detected in all individuals at 6 months post 2^nd^ dose and prior to the third vaccination suggesting the vaccine interval can influence class-switching to IgG4. Interestingly, unlike at 6-months post the 2^nd^ vaccine dose, where IgG1 responses had waned, IgG4 responses had increased. However, at 6-months post the 3^rd^ vaccine dose, all antibody isotypes had waned, including IgG4 (**Fig. 1C**).

Next, we examined the impact of hybrid immunity on IgG4 production, including infection prior to vaccination as well as a BTI post vaccination. A more heterogenous response was observed in individuals who had a SARS-CoV-2 infection prior to vaccination, with 3/8 of the volunteers producing Spike-specific IgG4 after the first vaccine dose, and all producing Spike-specific IgG4 after two doses (**Fig. 1D**). A breakthrough infection after three BNT162b2 vaccinations lead to a further increase in Spike-specific IgG4, reaching levels of only ∼5-fold below Spike-specific IgG1 in some individuals (**Fig. 1E**). Some individuals were subsequently vaccinated a fourth time with a bivalent WT/BA.1 mRNA vaccine, and this additional vaccination led to a further increase in Spike-specific IgG4 levels as well as total IgG, IgG1, IgG2 and IgG3 levels (**Fig. 1F**).

Together, these data demonstrate that Spike-specific IgG4 generated in response to mRNA vaccination is impacted by the interval and number of vaccinations, the time since vaccination, and an infection prior to/or after BNT162b2 vaccination.

### Spike-specific IgG4 is not detected in a longitudinal infection cohort

Several studies have reported a continued evolution of the antibody response post SARS-CoV-2 infection as demonstrated by increased somatic hypermutation of Spike-specific mAbs isolated 6-months post infection (Gaebler et al., 2021). To determine whether time from infection might influence the production of Spike- and/or nucleocapsid-specific IgG4 post infection we measured total IgG and IgG4 levels in a longitudinal infection cohort from Wave 1 (i.e. beginning March 2020) that was followed until 33 weeks post infection (**Table S1B** and **Fig. S1B**). Spike-specific IgG4 could not be detected in plasma from individuals experiencing mild (**Fig. 2A**) or severe (**Fig. 2B**) disease, even up to similar times post-infection as for the post-vaccination volunteers. These data suggest that the production of Spike-specific IgG4 is not an inherent feature to the Spike protein itself or the time post SARS-CoV-2 Spike exposure and that multiple exposures is important.

**Fig. 2.**
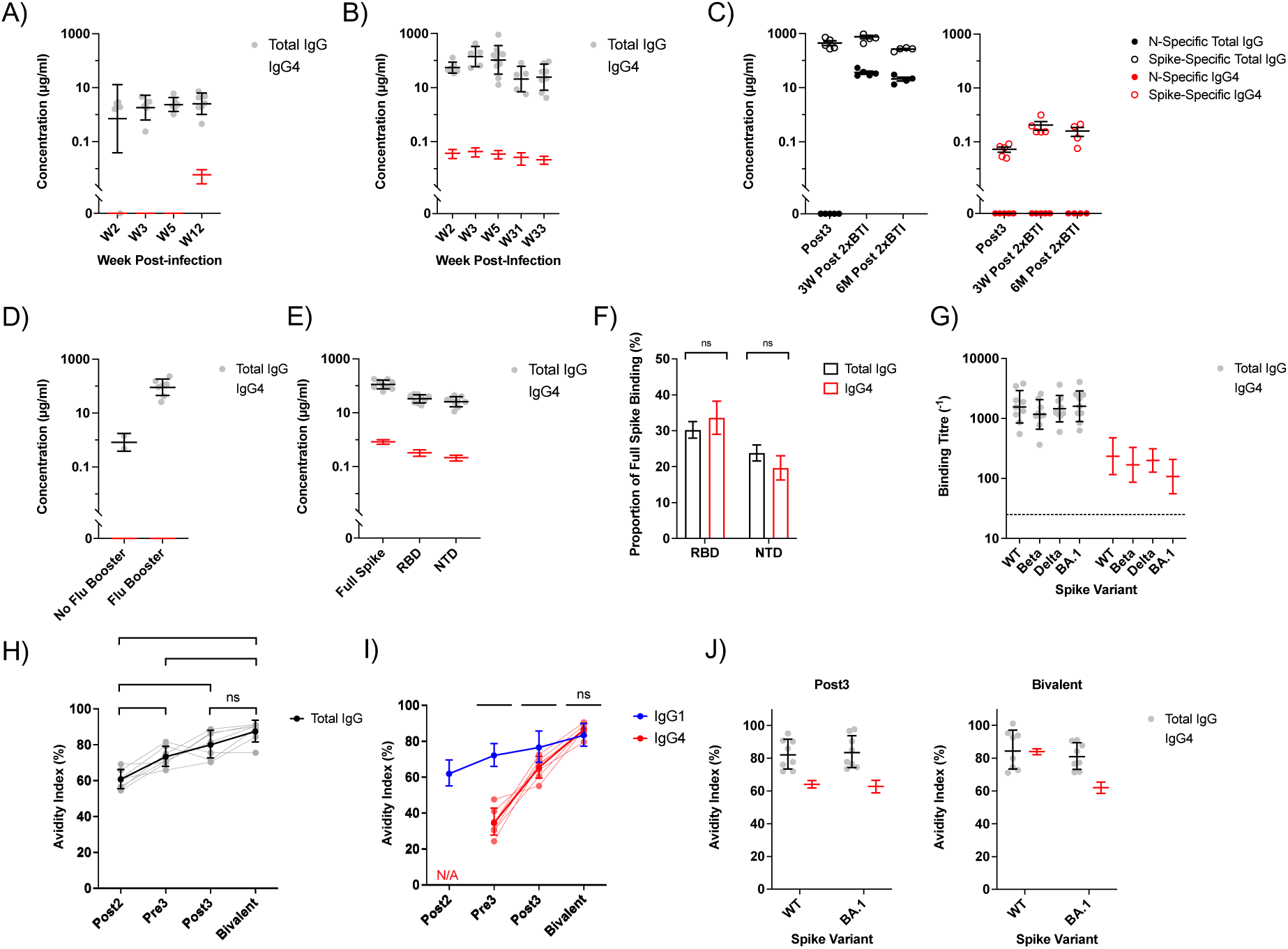
Specificity and avidity of IgG4 arising from BNT162b2 mRNA vaccination. Concentration of Spike-specific Total IgG (black/grey) and IgG4 (red/pink) in sera for **(A)** a mild infection group and **(B)** severe, ITU admission group. **(C)** Concentration of SARS-CoV-2 N-specific Total IgG (filled black circles), Spike-specific Total IgG (clear black circles), N-specific IgG4 (filled red circles), Spike-specific IgG4 (clear red circles), 2 weeks after 3^rd^ dose (Post3), 3 weeks after 2^nd^ breakthrough infection (3W Post 2xBTI) and 6 months after 2^nd^ breakthrough infection (6M Post 2xBTI). **(D)** Concentration of Haemagglutinin-specific Total IgG (black/grey) or IgG4 (red/pink) with or without influenza booster given with 12-week interval 3^rd^ Dose. **(E)** ELISA measurement of Total IgG (black/grey) and IgG4 (red/pink) targeted against WT SARS-CoV-2 Full Spike, Receptor-Binding Domain (RBD) and N-Terminal Domain (NTD) in 12-week interval Post3 samples. **(F)** RBD and NTD binding Total IgG (black) and IgG4 (red) as a proportion of binding to Full Spike. **(G)** Total IgG (black/grey) and IgG4 (red/pink) binding to WT, Beta, Delta and BA.1 Spike in 12-week interval Post3 samples. **(H)** Serum antibody avidity of Total IgG (black) binding to WT Spike in Post2, Pre3, Post3 and Bivalent samples from the Bivalent group. **(I)** As in (H), but for IgG1 (blue) and IgG4 (red). **(J)** Serum antibody avidity of Total IgG (black/grey) and IgG4 (red/pink) against WT and BA.1 recombinant Spikes for Post3 and Bivalent samples from the Bivalent group. In all panels, samples paired from an individual are in lighter colour, with darker coloured data showing geometric mean and standard deviation. Data representative of 3 technical replicates. ns=not significant, * p<0.05, ** p<0.01, *** p<0.001, **** p<0.0001

### IgG4 class-switch is Spike-specific

As the levels of Spike-specific IgG4 were boosted upon breakthrough infection, we next determined whether IgG4 production was restricted to Spike or if IgG4 to nucleocapsid (which is not present in the BNT162b2 vaccine) could be detected following breakthrough infection. After two breakthrough infections post-vaccination, there was a robust Spike-specific IgG4 response, which persisted for at least 6 months after the second infection (**Fig. 2C**). However, there was no nucleocapsid-specific IgG4 detected, even 6 months after a second breakthrough infection.

Some individuals also received an Influenza vaccine at the same time as their third BNT162b2 vaccination. We therefore investigated whether the immune environment generated in response to the mRNA vaccination might impact on IgG4 class-switching to influenza HA. No haemagglutinin-specific IgG4 were detected in any of these plasma samples (**Fig. 2D**). Combined, these data indicate that the class-switching to IgG4 is specific for SARS-CoV-2 Spike in the context of mRNA vaccination.

### Spike-specific IgG4 is distributed across RBD and NTD domains

We next investigated the domains on SARS-CoV-2 Spike targeted by the IgG4 response with a focus on the extended group. The two major Spike domains targeted by neutralising antibodies are the Receptor Binding Domain (RBD), which interacts with the host cell receptor Angiotensin Converting Enzyme 2 (ACE2) (Walls et al., 2020), and the N-Terminal Domain (NTD), which it’s function is less well defined (Jackson et al., 2022). Binding of IgG4 to both RBD and NTD could be detected by ELISA (**Fig. 2E**). When considering the proportion of IgG4 binding to each domain, 33.6% of the Spike-specific IgG4 were targeted against RBD, similar to 30.2% (SD 4.53) of the Total IgG; and 19.7% (SD 9.87) of the Spike-specific IgG4 was targeted against NTD, similar to 23.8% of the Total IgG (**Fig. 2F**) suggesting the Spike-specific IgG4 response is not skewed towards a specific Spike domain.

### Spike-specific IgG4 has similar variant binding breadth to total Spike-specific IgG

Previous studies have shown that repeated COVID-19 vaccination can lead to increased antibody binding and neutralisation breadth against newer SARS-CoV-2 variants of concern (Graham et al., 2022; Muecksch et al., 2022; Wratil et al., 2022). Given that Spike-specific IgG4 also emerges and increases following multiple mRNA vaccinations, we therefore measured breadth of the Spike-specific IgG1 and IgG4 responses by measuring variant Spike-binding breadth by ELISA. We focused on the Spike-specific Total IgG and IgG4 in the extended group following three BNT162b2 doses (**Fig. 2G**). Similar to the Total IgG binding, comparable levels of IgG4 binding were observed against the WT, Beta, Delta and BA.1 SARS-CoV-2 recombinant Spikes.

### Spike-specific IgG4 has a delayed avidity increase

Antibody avidity can also be enhanced by repeated antigen exposure, either through vaccination or infection (Graham et al., 2022; Wratil et al., 2022). Previous studies have associated the increased Spike-specific Total-IgG avidity with increasing levels of IgG4 (Irrgang et al., 2023). However, this study did not specifically measure avidity changes for Spike-specific IgG1 and IgG4 independently. Here, we first measured the avidity index against WT Spike of the Total IgG by comparing the area under the curve in ELISA, with and without an 8M Urea washing step (Graham et al., 2022). Similar to previously published results (Graham et al., 2022), the avidity of Spike-specific Total IgG increased with additional vaccine doses, as well as time post vaccination, up until the third dose where the avidity score reached a plateau (**Fig. 2H**). When considering the avidity of Spike-specific IgG1 and IgG4 independently, the IgG1 avidity mirrored the increasing avidity pattern observed for the Total IgG (**Fig. 2I**). In contrast, the avidity of Spike-specific IgG4 when first detected at 6-months post 2^nd^ vaccine was low (35.0%, SD 7.62) compared to the Total IgG (73.4%, SD 5.62) and to IgG1 (72.2%, SD 6.32) (**Fig. 2I**). However, the IgG4 avidity increased following a third BNT162b2 (65.3%, SD 5.84) with a smaller increase following a subsequent bivalent booster (86.4%, SD 3.44) to reach similar avidity levels to the Spike-specific IgG1 (83.4%, SD 6.23). Although the avidity against WT Spike became equal after a bivalent booster, the avidity against the BA.1 Spike remained lower for IgG4 (62.0%, SD 9.96) compared to IgG1 (81.4%, SD 8.13) (**Fig. 2J**) suggesting that affinity maturation of IgG4 against the BA.1 Spike is less. Overall, the avidity of Spike-specific IgG4 was delayed compared to IgG1 but plateaued at a similar level after three vaccine doses against the matched vaccine antigen.

### Serum antibody functional activity is associated with IgG1 levels, but not IgG4

We next investigated the functional effect of the increase in Spike-specific IgG4 on functional activity of polyclonal sera, including neutralisation, ADCC, ADCP and complement deposition. Given the largest increase in Spike-specific IgG4 was observed between the second and third dose of BNT162b2, we investigated the differences in functional activity between plasma from these time points in the extended group. Neutralisation activity is largely antibody Fc binding independent (Mozdzanowska et al., 2003), and as previously shown, there was a trend towards increasing plasma neutralising activity between the 2^nd^ and 3^rd^ dose of mRNA vaccine against WT (although this did not reach significance) consistent with the increase in Spike-specific Total IgG (**Fig. 3A**) (Graham et al., 2022). Both the IgG1 and IgG4 levels correlated with the ID_50_ values (**Fig. 3B**).

**Fig. 3.**
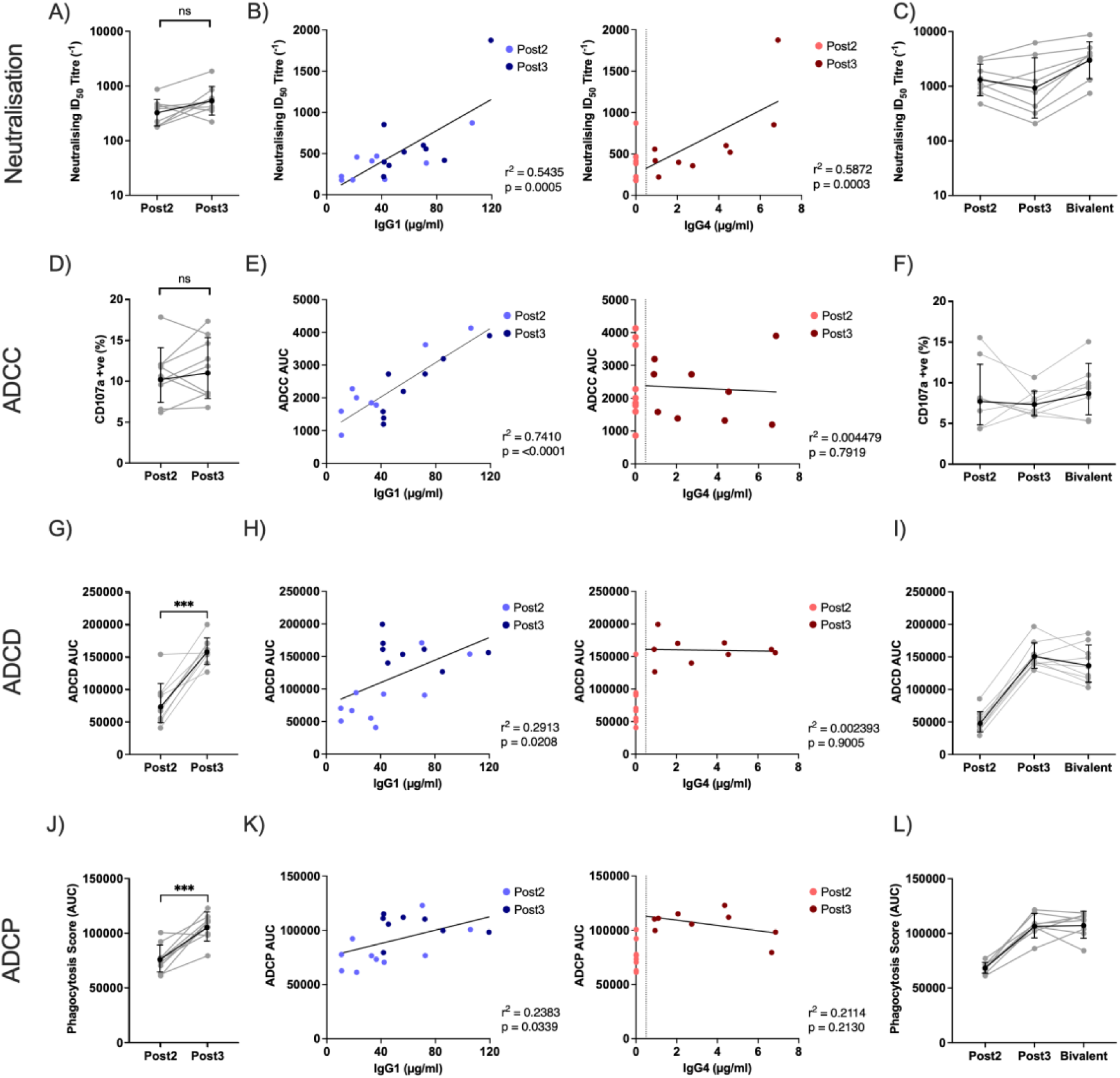
Functional activity of plasma with increasing Spike-specific IgG4. **(A)** Neutralising 50% Inhibitory Dose (ID_50_) titre against WT SARS-CoV-2 Spike pseudotyped virus for 12-week interval Post2 and Post3 samples. **(B)** Correlation between Neutralising ID50 and IgG1 (blue) or IgG4 (red) concentration, with lighter colour for Post2 and darker colour for Post3 samples. **(C)** Neutralising ID50 titre for Bivalent group Post2, Post3 and Bivalent samples. **(D, E & F)** as (A, B & C) but for Antibody Dependent Cellular Cytotoxicity (ADCC) assay, measuring surface CD107a on activated NK cells. **(G, H & I)** as (A, B & C) but for Antibody Dependent Complement Deposition (ADCD) assay, measuring deposited C3. **(J, K & L)** as (A, B & C) but for Antibody Dependent Cellular Phagocytosis (ADCP) assay, measuring THP-1 macrophage uptake of WT Spike-coated fluorescent beads incubated with plasma. For all correlations, best fit line drawn for all IgG1 data, and include data points with non-zero values of IgG4. The vertical dotted line represents the limit of IgG4 detection. Data representative of 3 technical replicates. ns=not significant, *** p<0.001

ADCC is dependent on CD16 expressed on NK cells, so we tested the ability of these plasma to stimulate an NK-92 cell line expressing CD16 to degranulate using CD107a expression as the readout (Bartsch et al., 2023). In our assay, there was no significant difference in NK cell degranulation between the 2^nd^ and 3^rd^ dose of mRNA vaccine at the highest dilution tested (1:50) (**Fig. 3D**), although there was notable heterogeneity between whether ADCC increased or decreased presumably due to the differing IgG1 and IgG4 levels. We therefore correlated the level of NK cell degranulation with concentration of Spike-specific IgG1 and IgG4. NK cell degranulation positively correlated with Spike-specific IgG1 levels, but no correlation was observed with Spike-specific IgG4 levels (**Fig. 3E**) or between the ratio of IgG1 to IgG4 (**Fig. S2A**). In the cohort receiving the bivalent vaccine, there was a similar trend between the 2^nd^ and 3^rd^ vaccine but the additional WT/BA.1 mRNA vaccination had no impact on ADCC activity despite small increases in Spike-specific IgG4 levels, suggesting a homeostatic limit to the amount of ADCC (**Fig. 3F**).

Next, we compared the ability of plasma collected following 2^nd^ and 3^rd^ dose of BNT162b2 vaccine to stimulate antibody-dependent complement deposition (ADCD) and antibody-dependent cellular phagocytosis (ADCP). As ADCD requires a complement source lacking antibodies to the SARS-CoV-2 Spike, we utilised Guinea Pig complement serum that was naïve to SARS-CoV-2 Spike (Polycarpou, 2023). Comparing ADCD activity in plasma post 2^nd^ and 3^rd^ dose showed a 3-fold increase in the level of complement deposition (**Fig. 3G**). There was a positive correlation between ADCD activity and IgG1 level, but no correlation was observed with IgG4 (**Fig. 3H**). ADCP activity was measured using differentiated THP-1 cells as the effector cells and fluorescent beads coated with recombinantly expressed SARS-CoV-2 Spike (Zohar et al., 2022). A 1.8-fold increase in ADCP activity was observed following the 3^rd^ mRNA vaccine (**Fig. 3J**). Similar to ADCD activity, there was a positive but weak correlation between ADCP activity and IgG1 level, but no correlation was observed with IgG4 (**Fig. 3K**). Following administration of the bivalent booster, ADCD and ADCP levels remained unchanged which is consistent with the smaller increase in Spike-specific IgG levels post bivalent booster (**Fig. 3I and 3L**).

A previous study reported a decrease in ADCD and ADCP following a third dose of BNT162b2 mRNA vaccine (Irrgang et al., 2023), however, in these experiments the serum input was normalised to the amount of Spike-specific Total IgG at each timepoint. When the ADCC, ADCD and ADCP activity measured in our experiments was normalised to the amount of Spike-specific Total IgG, we also observed a decrease in ADCD, and ADCP, as well as ADCC, for the Post3 samples compared to Post2 samples suggesting increasing IgG4 may limit effector function activity (**Fig. S2B**). No significant difference was observed for neutralisation. Nevertheless, given that there are increases in all IgG subclasses between Post2 and Post3 (**Fig. 1B**), the absolute ability of serum to activate these effector functions would be most physiologically relevant, rather than the relative functional ability.

In summary, between the 2^nd^ and 3^rd^ mRNA vaccine doses, where there is a 3.2-fold increase in IgG1 and an ∼100-fold increase in IgG4, we observe increased neutralisation, ADCD and ADCP in the plasma of individuals in the extended group. However, despite ADCC activity correlating with IgG1 concentration, ADCC activity in plasma does not increase significantly suggesting Spike-specific IgG4 impacts effector function activities differently.

### IgG4 maintains neutralisation activity, but loses Fc-dependent effector functions

Next, we investigated the specific contribution that IgG4 makes to the effector function activity measurements. Due to the technical challenges in separating the different subclasses of Spike-specific IgG from plasma we recombinantly expressed IgG1, IgG2, IgG3 and IgG4 versions of a panel of SARS-CoV-2 specific mAbs previously isolated from infected or vaccinated individuals (Graham et al., 2021; Seow et al., 2022). mAbs were selected based upon their ability to activate degranulation of CD16 expressing NK cells in the ADCC assay (**Fig. S3**). mAbs selected targeted several different epitopes on Spike. P008_60 is SD1 specific (Graham et al., 2021), P008_87 is a Class 3 RBD-specific mAb that binds an epitope on the outer face of RBD (Graham et al., 2021), VA14_1 and VA14_R39 are Class 4 RBD-specific mAbs that bind the interior face of RBD (Seow et al., 2022), and P008_99 is NTD-specific (Graham et al., 2021) (**Fig. S3A**). Note that none of the RBD Class 1 specific mAbs that bind the ACE2 receptor binding site were able to facilitate ADCC.

We first investigated the ability of immune complexes containing the different subclasses of IgG to bind to Fc receptors on the surface of various immune cells. NK cells express CD16 (FcγRIIIa), and incubation of mAb-immune complexes with NK cells demonstrated similar binding of IgG1 and IgG3 antibodies, but very low binding of the IgG2 and IgG4 subclasses (**Fig. S4A**). Raji cells expressing CD32 (FcγRIIa and FcγRIIb) (Cassel et al., 1993), bound immune complexes of all subclasses equally (**Fig. S4B**). THP-1 cells differentiated using PMA and IFNγ express both CD32 (FcγRII) and CD64 (FcγRI) (Fleit and Kobasiuk, 1991), and these showed weaker binding of IgG4 compared with other subclasses (**Fig. S4C**). Given that there was no difference in binding to Raji cells expressing CD32 alone, the difference in binding to the differentiated THP-1 cells is likely due to reduced binding between CD64 and the IgG4 Fc region.

Neutralisation of virus by IgG is largely antibody Fc-binding independent (Mozdzanowska et al., 2003), and indeed, class-switching each of these SARS-CoV-2 mAbs to IgG1, IgG2, IgG3 and IgG4 had no impact on their ability to neutralise ancestral SARS-CoV-2 pseudovirus infection of HeLa-ACE2 cells (**Fig. 4A**). Having demonstrated unchanged neutralisation activity, we next measured effector function activities of the class-switched antibodies. For all five mAbs, the IgG1 and IgG3 subclasses were able to stimulate CD16 expressing NK-92 cells to degranulate in the ADCC assay as measured by CD107a cell surface expression (Bartsch et al., 2023) (**Fig. 4B**). However, IgG2 and IgG4 subclasses were not able to activate degranulation of these cells, consistent with their inability to bind to the CD16 expressing NK-92 cell line (**Fig. S4A**).

**Fig. 4.**
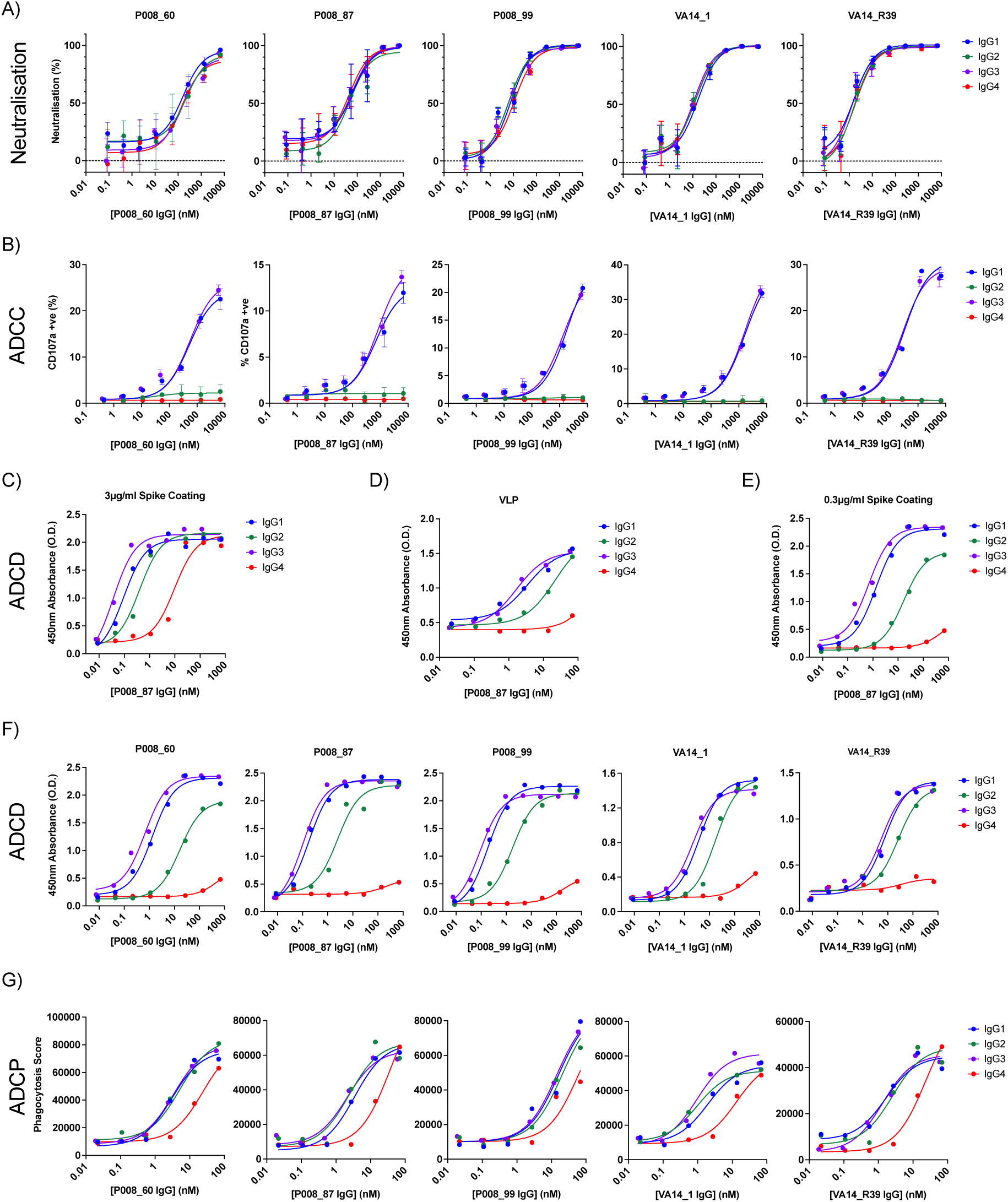
Functional activity of SARS-CoV-2 mAbs when expressed as different IgG subclasses. **(A)** Neutralisation curves for 5 monoclonal antibodies; P008_60, P008_87, P008_99, VA14_1, VA14_R39 class-switched to IgG1 (blue), IgG2 (green), IgG3 (purple) and IgG4 (red), against WT SARS-COV-2 Spike pseudotyped virus. **(B)** as in (A) but for ADCC measured as percentage of NK cells with surface-accessible CD107a. **(C)** ADCD assay for P008_87 as IgG1 (blue) or IgG4 (red) with 3 µg/ml WT Spike ELISA plate coating. **(D)** ADCD assay for P008_87 as IgG1 (blue) or IgG4 (red) against Spike Virus-like Particles (VLP) (filled circles) or Naked VLPs (clear circles). **(E)** ADCD assay for P008_87 as IgG1 (blue) or IgG4 (red) with 0.3 µg/ml WT Spike plate coating. **(F)** ADCD for P008_60, P008_87, P008_99, VA14_1, VA14_R39 class-switched to IgG1 (blue), IgG2 (green), IgG3 (purple) and IgG4 (red) with 0.3 µg/ml WT Spike plate coating. **(G)** as in (A) but for ADCP, measuring THP-1 macrophage uptake of WT Spike-coated fluorescent beads incubated with antibody. Data representative of 3 technical replicates.

In the complement activation assay, IgG1 and IgG3 versions of P008_87 were equally able to fix complement C3, and IgG2 and IgG4 displayed an 8-fold and 100-fold reduced complement fixation activity, respectively, when Spike was coated at 3 mg/mL (**Fig. 4C**). When the complement deposition assay was repeated with concentrated virus like particles (VLPs) expressing SARS-CoV-2 Spike coated on the ELISA plate, no ADCD activity of IgG4 could be detected (**Fig. 4D**) presumably due to a lower avidity of C3q binding. This hypothesis was confirmed, as ADCD activity of the IgG4 mAbs was also absent when the recombinant Spike was coated at a lower density of 0.3 mg/mL (**Fig. 4E**). A similar trend was observed for the remaining IgG4 mAbs on both concentrated VLPs (**Fig. S5**) and recombinant Spike coated at 0.3 mg/mL (**Fig. 4F**).

Finally, in the ADCP assay, equal phagocytic activity was observed for the IgG1, IgG2 and IgG3 subclasses of each mAb, but a 5 to 15-fold reduction in the phagocytic activity of IgG4 which is consistent with the reduced binding to the differentiated THP-1 cells (**Fig. 4G**). Overall, the IgG4 subclasses were unable to facilitate ADCC and had reduced ADCD and ADCP activity.

### Non-competing IgG4 is functionally silent, not inhibitory

Antibodies function *in vivo* within a polyclonal mix of antigen-specific antibodies of different isotypes and subclasses. To determine the functional impact of Spike-specific IgG4 on overall antibody effector function of plasma, we next investigated how the mAbs of different IgG subclasses, either targeting the same epitope or non-competing epitopes, would interact with one another in effector function assays.

First, we investigated the effect of the presence of IgG4 targeting the same epitope as an IgG1 with potent ADCC activity. We maintained a constant level of the IgG1 version of the mAb (either P008_87 or VA14_R39), prior to titrating in either additional IgG1 or IgG4 of the same mAb. Adding in additional IgG1 of the mAb led to an increased NK cell degranulation once the concentration was greater than the baseline level of mAb (**Fig. 5A**). However, addition of the IgG4 version of the mAb binding to the same epitope led to inhibition of degranulation, likely due to competition for binding to a shared epitope on Spike. The same trend was observed for both P008_87 and VA14_R39. However, this inhibition of ADCC activity was not observed when an IgG4 targeting a non-competing epitope was added (**Fig. 5B**). Again, a constant level of IgG1 antibody was maintained, but either an additional IgG1 or IgG4 of a non-competing mAb was added. Addition of IgG1 similarly led to additional NK cell degranulation. However, addition of IgG4 as a non-competing antibody had no effect on overall degranulation. A similar experimental approach was used to study the direct impact of Spike-specific IgG4 on ADCD by a potent IgG1 mAb. When IgG1 was added to a constant level of IgG1 targeting the same epitope (either P008_87 or VA14_R39), an increase in ADCD was observed when the IgG4 version of this mAb was added a decrease in ADCD indicating an inhibitory effect (**Fig. 5C**). However, addition of an IgG1 or IgG4 targeting a different Spike epitope, showed no decrease in ADCD activity (**Fig. 5D**). Overall, the presence of Spike specific IgG4 could potentially decrease effector function activity (both ADCC and ADCP) through preventing the binding of IgG subclasses that are potent activators of effector functions, including IgG1.

**Fig. 5.**
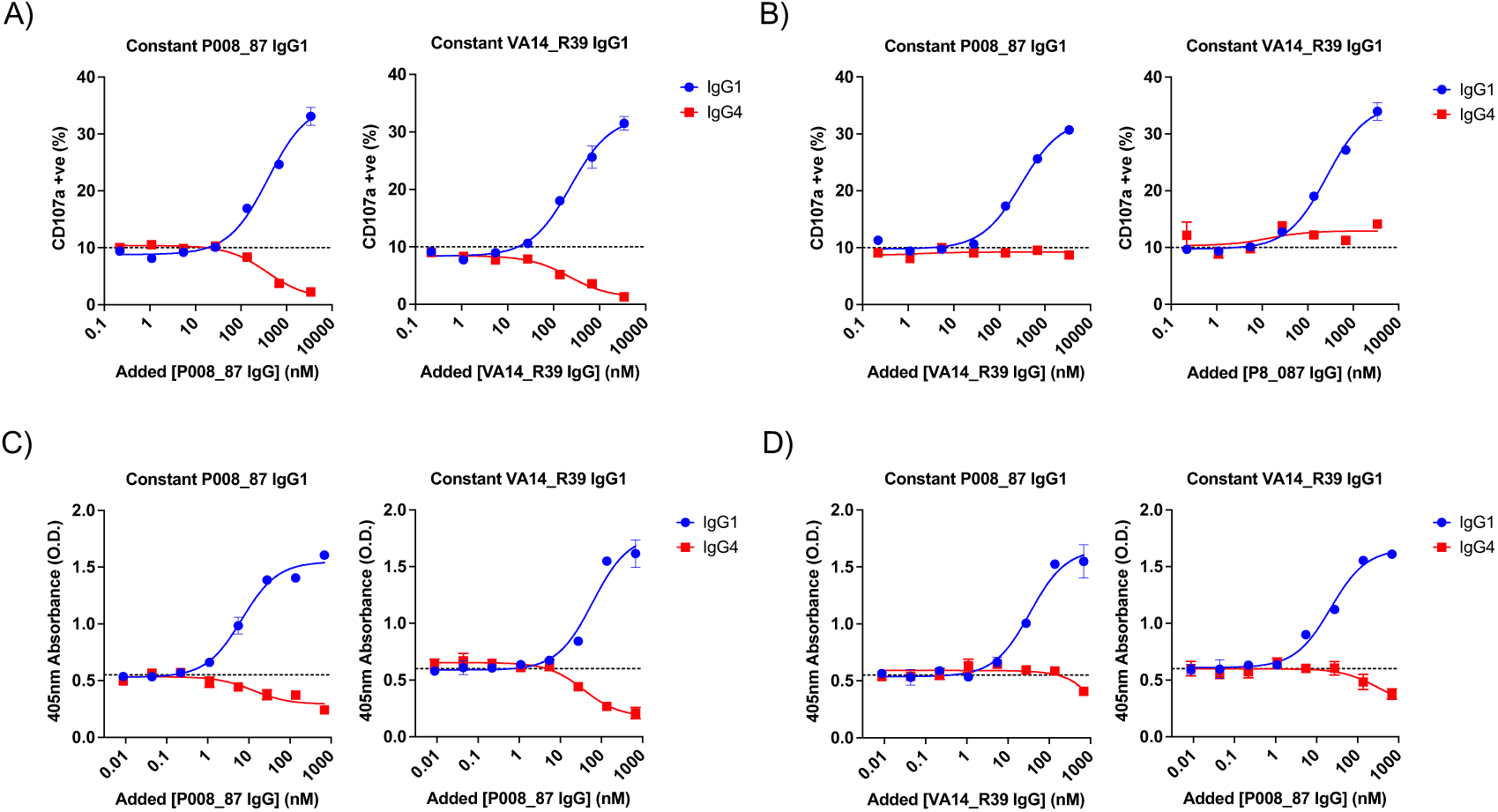
Effector function competition assays between IgG1 and IgG4. **(A)** ADCC competition assay, with constant level of either P008_87 IgG1 or VA14_R39 IgG1 to give 10% NK cells with detectable surface CD107a, incubated with increasing amounts of the same antibody as an IgG1 (blue) or IgG4 (red). **(B)** ADCC competition assay with constant level of either P008_87 IgG1 or VA14_R39 IgG1 to give 10% of NK cells surface CD107a, incubated with increasing amounts of the other non-competitively binding antibody. **(C)** ADCD competition assay with constant level of either P008_87 IgG1 or VA14_R39 IgG1, incubated with increasing amounts of the same antibody as IgG1 (blue) or IgG4 (red). **(D)** ADCD competition assay with constant level of P008_87 IgG1 or VA14_R39 IgG1, incubated with increasing amounts of the other non-competitively binding antibody. Data representative of 2 technical replicates.

To determine whether such an effect might be seen *in vivo* in the context of a polyclonal antibody response, we added a cocktail of the five IgG4 mAbs targeting different Spike epitopes to plasma collected after two mRNA doses where Spike-specific IgG4 level is very low and measured the impact on ADCD and ADCC (**Fig. S6A-B**). Pooled IgG4 or IgG1 was added at a combined concentration of 1 mg/mL, which greatly exceeds the maximum level of Spike-specific IgG4 measured in plasma. Addition of pooled Spike-specific IgG1 mAbs was able to increase the level of ADCD and ADCC (**Fig. S6A** and **S6B**). However, although the pooled IgG4 was able to compete the activity of IgG1 mAb P008_99 (**Fig. S6A** and **S6B**), no difference in ADCD or ADCC activity was observed for the polyclonal plasma. This difference may be due to the recombinant IgG4s only targeting a proportion of the potential Spike epitopes targeted by the functional IgG1 and IgG3 in the polyclonal plasma. These data suggest that the interplay between IgG1 and IgG4 within the context of the polyclonal sera where there is a wider array of neutralising and non-neutralising epitopes targeted on Spike, is more complex.

## Discussion

This study supports previous observations that the COVID-19 mRNA based vaccine, BNT162b2, can lead to a Spike-specific IgG4 response following multiple exposures (Akhtar et al., 2023; Buhre et al., 2022; Gelderloos et al., 2024; Irrgang et al., 2023). This study extends these observations further through additional understanding of the factors impacting switch to IgG4 following BNT162b2 vaccination and the functional activity and potential consequences of high levels of Spike-specific IgG4.

Comparing the kinetics and magnitude of Spike-specific IgG4 production in individuals with varying vaccination histories enabled us to identify additional factors that impact IgG4 class-switching in the context of mRNA vaccination. The interval between first and second BNT162b2 dose proved important for the switch to Spike-specific IgG4 as demonstrated by an elevated level of IgG4 in the extended (12-week interval) vs short (3-week interval) group at 6-months post second vaccine dose. However, a third BNT162b2 dose resulted in boosting to reach similar levels of Spike-specific IgG4 in both the short and extended groups. This data suggests that vaccine interval alone is not sufficient to explain the previously reported difference in production of Spike-specific IgG4 between BNT162b2 administered at a 3-week interval and AZD1222 administered at a 12-week interval, where no Spike-specific IgG4 was detected, but the difference must also be related to the vaccine type and formulation (Hartley et al., 2024). We observed the level of Spike-specific IgG4 to further increase following additional mRNA vaccination similar to Gelderloos *et al* who showed that the IgG4 level can continue to increase after up to 5-doses of mRNA vaccine in older adults (Gelderloos et al., 2024). Spike antigen exposure through breakthrough infection also increased IgG4 Spike levels. Whether these increases are due to continued evolution of already switched IgG4 or due to new class-switching events needs to be investigated. However, the class-switch to IgG4 is specific for SARS-CoV-2 Spike in the form of an mRNA-based immunogen as Nucleoprotein-specific IgG4 was not detected following two SARS-CoV-2 breakthrough infections, and HA-specific IgG4 was not detected in individuals co-immunised with the influenza vaccine at the time of mRNA booster vaccination. These results indicate there may be specific conditions created by the mRNA vaccine that make B cells more prone to induce class switch recombination (CSR) to distal subclasses. Investigation of class-switching to antigen-specific IgG4 for other mRNA-based vaccines is required.

It has been proposed that long germinal centre reactions observed following mRNA vaccination, where there is a prolonged presence of the Spike antigen in the lymph node (Kim et al., 2022; Roltgen et al., 2022; Turner et al., 2021), might explain the high-level of Spike-specific IgG4 (Irrgang et al., 2023). Prolonged antigen exposure has been proposed as the mechanism behind generation of IgG4 responses to honey bee venom observed in bee keepers (Aalberse et al., 1983). However, prolonged antigen exposure through repeated vaccination with tetanus vaccines is not sufficient to generate a vaccine specific IgG4 response (Unger et al., 2018). Furthermore, despite continued maturation of the Spike-specific IgG response being observed for at least 6-months post SARS-CoV-2 infection, presumably due to extended antigen exposure (Gaebler et al., 2021), no Spike-specific IgG4 was detected in our longitudinal infection cohort, including both mild and severe disease, up to 33 weeks post infection. These observations highlight that repeated or prolonged antigen exposure is not sufficient to facilitate IgG4 switch.

The distribution in IgG4-binding to the RBD and NTD domains was similar to that observed for total IgG and IgG1 suggesting there is no epitope bias in the subclass switching. As the IgG4 shows a similar breadth of variant Spike binding it also suggests that the IgG4 B cells could be arising from re-activation and switching of existing Spike-reactive B cells rather than a *de novo* response. Additionally, SARS-CoV-2 infection prior to BNT162b2 vaccination was observed to accelerate the appearance of Spike-specific IgG4, with IgG4 first being detected at low levels from 3 weeks post 1^st^ BNT162b2 dose in 3/8 individuals and in all (8/8) individuals from 3 weeks post 2^nd^ dose. This more rapid appearance suggests that IgG subclasses generated in response to SARS-CoV-2 infection can be class-switched to IgG4 upon mRNA vaccination indicative of reactivation of existing B cell clones rather than generation of a *de novo* response. However, it is noteworthy that the avidity of the Spike-specific IgG4 lags behind that of the Total-IgG, particularly against the BA.1 variant Spike. In contrast to the former observation, this may indicate that IgG4 production does not primarily arise from the class-switching of high avidity IgG subclasses. Alternatively, the lower avidity could also arise due to the ability of IgG4 to undergo Fab arm exchange leading to monovalent antibody binding with reduced avidity (van der Neut Kolfschoten et al., 2007). Further investigations into the clonal origin of IgG4 in relation to other antibody classes and IgG subclasses over time is required to understand the origin of IgG4 further.

Antibody effector functions have been shown to be important in the therapeutic and prophylactic activity of SARS-CoV-2 mAbs in animal models (Gorman et al., 2021; Zhang et al., 2023). A significant concern with the elicitation of Spike-specific IgG4 is the reported anti-inflammatory properties arising from reduced affinity to Fcγ receptors (Pillai, 2023; Vidarsson et al., 2014). Importantly, despite the production of Spike-specific IgG4, repeated mRNA vaccine boosting has remained protective from severe COVID-19 (Lee et al., 2023). The consequence of high levels of Spike-specific IgG4 on effector function activity is difficult to tease apart due to the polyclonal nature of plasma, and the varying proportions and amounts of Spike-specific IgG1, IgG2, IgG3 and IgG4 both at different timepoints and between individuals, as well as the lack of animal models that produce IgG4 (Olivieri and Gambon Deza, 2018). By isotype-switching previously characterised Spike-specific mAbs, we have been able to investigate the functional implications of elevated levels of Spike-specific IgG4 more directly. As expected, antibody class-switching had no impact on the ability of mAbs to neutralise SARS-CoV-2 which is consistent with the increasing plasma neutralising activity observed as Spike-specific Total IgG increases. Spike-specific IgG1 and IgG3 were able to facilitate ADCC, ADCD and ADCP equally. Spike-specific IgG2 lost ADCC activity, retained ADCP activity and was a weak activator of ADCD. Spike-specific IgG4 had the lowest levels of effector function activity, showing no ADCC and ADCD activity (at low antigen densities) and had a 10-fold reduction in ADCP. Using mAbs expressed as different subclasses in ADCC and ADCD competition assays, we showed that in the context of viral infection the Spike-specific IgG4 is functionally silent unless there is direct binding competition with a functionally active IgG1. In this scenario the IgG4 reduces effector function activity by preventing the binding of IgG1 to the Spike antigen. In the context of allergy, increased titres of allergen specific IgG4 has been shown to reduce hypersensitivity through blocking the activity of IgE (Santos et al., 2015) and IgG4 in the context of mRNA vaccination may have a comparative function in reducing inflammatory responses whilst maintaining the ability to neutralise free virus. Indeed, this blocking nature of IgG4 makes it a commonly used format for therapeutic monoclonal antibodies due to its limited ability to induce inflammatory responses (Rispens and Huijbers, 2023). Despite the unusual presence of Spike-specific IgG4, the relatively low-level present compared to the more functionally active IgG1 suggests the inhibitory effect arising from steric occlusion could be out-competed for the limited antigenic sites on the surface of a virion or infected cell or the lower avidity of IgG4 binding. Furthermore, in the context of SARS-CoV-2 infection, antibodies against viral proteins other than Spike can also mediate effector functions such as ADCC (Fielding et al., 2022). Therefore, the non-stimulatory effect of Spike-specific IgG4 may be hidden in more complex scenarios, such as polyclonal mixes. Indeed, we were unable to inhibit ADCD or ADCC activity in polyclonal sera through addition of a cocktail of Spike-specific IgG4s at high concentrations (up to 1 mg/mL).

In summary, we show that factors including vaccine interval and frequency, hybrid immunity and breakthrough infection all enhance the production of Spike-specific IgG4 following BNT162b2 mRNA vaccination. Through generation of Spike-specific mAbs of each IgG subclass, we showed that the IgG4 can inhibit ADCC and ADCD activity of potently activating IgG1 through direct binding competition. However, the polyclonal nature of plasma and the lower avidity of IgG4 binding observed suggest that the impact on overall effector function activity is likely to be minimal, and will be heavily dependent on the epitope specificity of the IgG4 antibodies produced. Further research is needed to understand the factors that trigger IgG4 class switching in the context of mRNA vaccination and the origin of the antigen-specific IgG4 antibodies.

## Materials and methods

### Volunteer Samples/Ethics

Demographics of the vaccinated and infected volunteer samples are shown in **Supplementary Table 1**. Samples were collected according to the timeline shown in **Supplementary Fig. S1**. This study used human samples collected with written consent as part of a study entitled “Antibody responses following COVID-19 vaccination.” Ethical approval was obtained from the King’s College London Infectious Diseases Biobank (IBD) (KDJF-110121) under the terms of the IDB’s ethics permission (REC reference: 19/SC/0232) granted by the South Central Hampshire B Research Ethics Committee in 2019 and London Bridge Research Ethics Committee (reference: REC14/LO/1699). Collection of surplus serum samples at St Thomas Hospital, London, was approved by South Central-Hampshire B REC (20/SC/0310).

### Bacterial Strains and Cell Culture

SARS-CoV-2 pseudotypes were produced by transfection of HEK293T/17 cells (ATCC CRL-11268) and neutralisation activity assayed using HeLa cells stably expressing ACE2 (kind gift from Dr James Voss, Scripps Research, CA). Monoclonal antibodies were expressed in HEK293 Freestyle (HEK293F) cells (Thermofisher Scientific). NK-92 cells (obtained from ATCC, CRL-2407) transduced with high affinity (Valine 158) human CD16, THP-1 (TIB-202) and Raji (CCL-86) cells were obtained from ATCC. HEK293T/17 cells and HeLa-ACE2 cells were maintained in Dulbecco’s Modified Eagle Medium supplemented with GlutaMAX, 10% Foetal Calf Serum and 1% Penicillin/Streptomycin (DMEM-C). HEK293F cells were maintained in Freestyle media. THP-1 and Raji cells were maintained in Roswell Park Memorial Institute (RPMI) 1640 supplemented with GlutaMAX, 10% Foetal Calf Serum (RPMI-C). NK-92 cells expressing CD16 were maintained in Minimum Essential Medium α (MEM α) supplemented with L-glutamine, nucleosides, 12.5% Foetal Calf Serum, 12.5% Horse Serum, 20mM HEPES, 0.2mM Myo-inositol, 0.02mM Folic Acid, 0.1mM 2-mercaptoethanol and 50IU/ml Interleukin (IL)-2. Bacterial transformations for class switching were performed with NEB Stable Competent *E.coli*.

### Protein Expression and Purification

Recombinant Spike, RBD and NTD (residues 1-310) were expressed and purified as previously described (Rosa et al., 2021; Seow et al., 2020). Recombinant SARS-CoV-2 WT Nucleocapsid (N protein) was a kind gift from Dr Leo James, Laboratory of Molecular Biology, Cambridge. Recombinant Spike trimers were engineered from full-length Spike expression plasmids (pcDNA3.1 plasmid) using a molecular cloning approach. The furin cleavage site (RRAR) was replaced with a flexible linker (GGGG), and the K986 and V987 residues in the S2 subunit were mutated to prolines using site-directed mutagenesis. The resulting sequences were truncated at amino acid position 1138 and subcloned into pHLSec (Aricescu et al., 2006) via Gibson assembly, positioning them upstream of a flexible linker (GSGG), T4 foldon trimerisation domain, and incorporating Avi and His tags.

Biotinylated Spike was expressed as previously described (Graham et al., 2021). Biotinylated Spike was expressed in 1L of HEK293F cells (Invitrogen) at a density of 1.5 × 10^6^ cells/mL. To achieve *in vivo* biotinylation, 480µg of each plasmid was co-transfected with 120µg of BirA (Howarth et al., 2008) and 12mg PEI-Max (1 mg/mL solution, Polysciences) in the presence of 200 µM biotin (final concentration). The supernatant was harvested after 7 days and purified using immobilised metal affinity chromatography and size-exclusion chromatography. Complete biotinylation was confirmed via depletion of protein using avidin beads.

Influenza HA haemagluttinin A/California 04/2009 was cloned into the pHLSec vector containing a 6xHIS tag, and expressed and purified in the same manner as for SARS-CoV-2 Spike.

### IgG subclass cloning and expression

SARS-CoV-2 monoclonal antibodies were isolated and characterised previously (Graham et al., 2022; Seow et al., 2022). Heavy chains were PCR amplified using previously described primers and conditions (Graham et al., 2022; Seow et al., 2022; Tiller et al., 2008), and cloned into human IgG heavy chains of IgG2 (Addgene AbVec2.0-IGHG2 #99576), IgG3 (AbVec2.0-IGHG3 #99577) and IgG4 (AbVec2.0-IGHG4 #99578) using the Gibson Assembly Master Mix (NEB) following the manufacturer’s protocol. AbVec2.0-IGHG2, AbVec2.0-IGHG3 and AbVec2.0-IGHG4 were a gift from Hedda Wardemann (Addgene plasmid #99576, #99577 and #99578; RRID:Addgene_99576, RRID:Addgene_99577 and RRID:Addgene_99578).

Ab heavy and light plasmids were co-transfected at a 1:1 ratio into HEK-293F cells (Thermofisher) using PEI Max (1 mg/mL, Polysciences, Inc.) at a 3:1 ratio (PEI Max:DNA). Ab supernatants were harvested five days following transfection, filtered and purified using protein G affinity chromatography following the manufacturer’s protocol (GE Healthcare).

### Pseudovirus production

HEK293T/17 cells were seeded in a 10cm dish at a density of 3×10^5^cells/mL. Following overnight culture, cells were co-transfected using 90 µg PEI-Max (1 mg/mL, Polysciences) with 15 µg HIV-luciferase plasmid, 10 µg HIV 8.91 gag/pol plasmid, and 5 µg SARS-CoV-2 Spike protein plasmid (Zufferey et al., 1997). Transfected cells were incubated for 72 h at 37°C, and virus was harvested, sterile filtered, and stored at −80°C until required.

### VLP production and quantification

HEK293T/17 cells were seeded in a 10cm dish at a density of 3×10^5^ cells/mL. Cells were transfected using 90µL PEI with the following plasmids: 2.5µg WT SARS-CoV-2 Spike plasmid, 8µg HIV gag/pol mScarlet plasmid, 2µg HIV 8.91 gag/pol plasmid, and 10µg HIV-luciferase plasmid. The supernatant containing the VLPs was collected 72h post-transfection, clarified by centrifugation at 3000×g for 15 min and passed through a 0.45µm filter. The clarified supernatant was then layered onto a sucrose cushion (20% sucrose, 50 mM Tris-HCl, 0.5 mM EDTA, and 100 mM NaCl in PBS, pH 7.4) and centrifuged at 10000×g for 4h at 4°C. The supernatant was gently aspirated and the concentrated VLPs were resuspended in PBS. The samples were then stored at −80°C until use. Infectivity of VLPs was measured using HeLa ACE2 cells.

### Antibody Subclass ELISA

High-binding ELISA plates (Corning, 3690) were coated with antigen (N protein, S glycoprotein, RBD, NTD or Influenza HA) at 3 μg/mL (25 μl per well) in phosphate-buffered serum (PBS) overnight at 4 °C. Wells were washed with PBS-T (PBS with 0.05% Tween-20) and then blocked with 100 μl of 5% milk in PBS-T for 1 h at room temperature. The wells were emptied, and serial dilutions of plasma (heat-inactivated at 56°C for 30 min) were added and incubated for 1 h at room temperature. Dilutions of a known monoclonal antibody standard, and serum antibody standard ERM-DA470k (Sigma) (Wilson et al., 2013) were used for quantification. Wells were washed with PBS-T. Secondary antibodies were added (1:1000 dilution in 5% No Fat Milk in PBS-T) and incubated for 1 h at room temperature, including goat-anti-human Total IgG-alkaline phosphatase (AP) (Jackson Immunoresearch, 109-055-098), mouse-anti-human IgG1-horseradish peroxidase (HRP) (Invitrogen A10648), mouse-anti-human IgG2-HRP (Invitrogen MH1722), mouse-anti-human IgG3-HRP (Southern Biotech 9210-05), mouse-anti-human IgG4-HRP (Invitrogen MH174225)). Wells were washed with PBS-T and either AP substrate (Sigma) was added and read at 405 nm (AP) or one-step 3,3′,5,5′-tetramethylbenzidine (TMB) substrate (Thermo Fisher Scientific) was added and quenched with 0.5 M H_2_S0_4_ before reading at 450 nm (HRP). Quantification was conducted by comparing dilutions to reach the same EC_50_ value calculated by GraphPad Prism.

### Avidity Assay

The avidity ELISA was carried out as described above except for one additional step. After incubation of plasma, one half of the plate was incubated with 8M Urea and the other half incubated with PBS for 15 min before washing 5-times with PBS-T. The area under the curve was determined in Prism (Log dilution). The avidity index was calculated using the following formula:

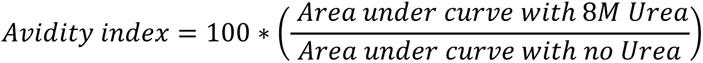

### Immune Complex Binding Assay

NK-92 cells expressing human CD16, THP-1 cells and Raji cells were plated at 1.0×10^5^cells/well on high-binding full-well 96 well plates. Immune complexes were formed by incubating 25µl of serially diluted mAb in RPMI-C (at concentrations stated), with WT Spike protein (25µl at 3.0 µg/ml) for 1h. Immune complexes were then incubated with each cell type for 1h. Cells were then washed with PBS-T, and blocked with Fc Block (BD Biosciences) for 1h. Following a final was in PBS-T, binding to cells was measured using the same secondary antibodies as for ELISA above.

### Neutralisation Assay

Serial dilutions of plasma samples (heat inactivated at 56°C for 30mins) or monoclonal antibody were prepared in DMEM-C (to total 25μL) and incubated with pseudotyped virus (25μL) for 1h at 37°C in half-area 96-well plates. HeLa ACE2 cells were diluted to a concentration of 5×10^5^cells/mL and added to each well (25µL/well), and plates incubated at 37°C for 72h. Infection levels were quantified by lysing the cells and measuring luciferase activity using the Bright-Glo Luciferase Assay Kit (Promega) on a Victor X3 multilabel plate reader (PerkinElmer).

### Antibody Dependent Cellular Cytotoxicity (ADCC) Assay

ADCC assay was adapted from Bartsch *et al* (Bartsch et al., 2023). High-binding full-well ELISA plates were coated with Spike protein at 3 μg/mL (50 μl per well) in PBS for 2h at 37 °C. Wells were washed with PBS and then blocked with 100 μl of 5% bovine serum albumin in PBS overnight at 4°C. The plates were then washed with PBS, prior to addition of serial dilutions of heat inactivated plasma samples or monoclonal antibody diluted in DMEM-C and incubated at room temperature for 1h. NK-92 CD16 cells were added (1.0×10^5^cells/well) in media supplemented with an increased IL-2 concentration (100 IU/ml), Golgistop protein transport inhibitor (BD Biosciences #554724, 5µl per 1.0×10^6^cells) and PerCP/Cyanine5.5 anti-human CD107a antibody (Biolegend Clone H4A3). Plates were incubated at 37°C, 5% CO_2_ for 6h. NK-92 cells were transferred to 75mm Polystyrene Tubes, washed with PBS and fixed by addition of 4% paraformaldehyde. Level of CD107a surface expression was measured by flow cytometry using a BD Canto II.

### Antibody Dependent Complement Deposition (ADCD) Assay

ADCD assay was adapted from Ref (Polycarpou, 2023). High-binding ELISA plates (Corning, 3690) were coated with Spike protein at 3 μg/mL or 0.3 μg/mL (25 μl per well) in phosphate-buffered serum (PBS) overnight at 4 °C. For VLP assays, purified VLPs at 4×10^6^ infectious units/well were incubated at room temperature for 2h. Wells were washed with PBS-T (PBS with 0.05% Tween-20) and then blocked with 100 μl of 5% milk in PBS-T for 1h at room temperature. Serial dilutions of heat-inactivated plasma or mAbs (25µl in PBS) were added and incubated for 1h at room temperature, followed by washing with PBS-T. Guinea Pig Complement Serum (Sigma S1639) was diluted to 1:100 (as per the complement activity level) in HEPES Buffered Saline (0.01 M HEPES, 0.15 M NaCl, 135 nM CaCl_2_, 1 mM MgCl_2_, adjusted to pH 7.4) as barbitone buffers are not available (Zelek et al., 2018). Diluted complement was added to the wells, alongside a heat inactivated control plasma (56°C for 30mins), and plate was incubated for 1h. Wells were washed with PBS-T, prior to addition of Goat Anti-Guinea Pig Complement C3 polyclonal (1:250 in 5% milk/PBS-T, MP Biomedicals 0855371) for 1h at room temperature. Wells were washed with PBS-T, and incubated with HRP-conjugated Mouse Anti-Goat IgG (1:500, Santa Cruz Biotechnology sc-2354). Wells were then washed and TMB substrate added which was quenched with 0.5 M H_2_S0_4_ before reading at 450 nm (HRP).

### Antibody Dependent Cellular Phagocytosis (ADCP) Assay

The ADCP assay was based on previously published protocol (Zohar et al., 2022). Briefly, biotinylated SARS-CoV-2 Spike (5μL at 1 mg/mL) was incubated with 10µl of washed yellow-green fluorescent NeutrAvidin-Labeled Microspheres (480-505nm, 1.0μm size, Thermofisher), and incubated at 37 °C for 2h. Beads were washed twice (0.1% PBS-BSA) by centrifugation (16000xg, 5min) at room temperature. The antigen-coupled fluorescent beads were resuspended in 0.1% PBS-BSA (1:500 by volume). 2.5µl of this bead suspension was incubated with 5µl heat inactivated plasma or antibody dilutions at room temperature for 1h. THP-1 cells (25,000/well) and bead/plasma mixes were added to 24 well plates and incubated for 16h at 37°C, 5% CO_2_. Cells were washed and bead uptake measured by flow cytometry using a BD FACS Canto II. Phagocytosis scores were determined by first plotting the antibody concentration against the geometric mean fluorescence intensity of the bead positive THP-1 cells multiplied by the percent of bead positive THP-1 cells. These curves were used to calculate the AUC using GraphPad Prism.

### IgG Subclass ADCC and ADCD Competition Assays

For competition assays, IgG1 of P008_87 or VA14_R39 were titrated to either 10% NK-92 cell degranulation activity or 25% complement C3 detection (0.5 at 450nm absorbance). Further P008_87 or VA14_R39 of IgG1 or IgG4 subclass was then added into this antibody mixture at different serial dilutions. These antibody mixes were then used in the ADCC and ADCD assays as described above.

### Plasma Spiked IgG4 Assay

P008_60, P008-87, P008_99, VA14_1 and VA14_R39 IgG1 and IgG4 were pooled together for final concentrations of 1 mg/ml each. The IgG4 mixture was serially diluted and 25uL added to plasma samples (25µl of EC_80_ values calculated from a prior ADCD assay (roughly 1:100 dilution)). P008_99 IgG1 mAb was used at 1 µg/ml as an ADCD and ADCC positive control.

## Acknowledgements

We thank Wendy Barclay for providing the Beta, Delta, BA.1 Spike expression plasmids and James Voss for providing the HeLa-ACE2 cells. We thank Kareena Kumar for technical assistance.

This work was funded by; MRC project grant ([MR/X009041/1] to KJD), MRC Genotype-to-Phenotype UK National Virology Consortium ([MR/W005611/1] and [MR/Y004205/1] to KJD), and Wellcome funded consortium - Genotype-to-Phenotype Global ([226141/Z/22/Z] to KJD). Fondation Dormeur, Vaduz for funding equipment to KJD. KJD was supported by the Medical Research Foundation Emerging Leaders Prize 2021. RJS was supported by the MRC (MR/S00971X/1) and Wellcome Trust (226615/Z/22/Z).

## Notes

### Competing Interest Statement

The authors have declared no competing interest.

## References

Aalberse, R.C., van der Gaag, R., and van Leeuwen, J. (1983). Serologic aspects of IgG4 antibodies. I. Prolonged immunization results in an IgG4-restricted response. J Immunol 130, 722–726.

Akhtar, M., Islam, M.R., Khaton, F., Soltana, U.H., Jafrin, S.A., Rahman, S.I.A., Tauheed, I., Ahmed, T., Khan, II, Akter, A., et al. (2023). Appearance of tolerance-induction and non-inflammatory SARS-CoV-2 spike-specific IgG4 antibodies after COVID-19 booster vaccinations. Front Immunol 14, 1309997.

Aricescu, A.R., Lu, W., and Jones, E.Y. (2006). A time- and cost-efficient system for high-level protein production in mammalian cells. Acta Crystallogr D Biol Crystallogr 62, 1243–1250.

Baden, L.R., El Sahly, H.M., Essink, B., Kotloff, K., Frey, S., Novak, R., Diemert, D., Spector, S.A., Rouphael, N., Creech, C.B., et al. (2021). Efficacy and Safety of the mRNA-1273 SARS-CoV-2 Vaccine. N Engl J Med 384, 403–416.

Bartsch, Y.C., Cizmeci, D., Kang, J., Gao, H., Shi, W., Chandrashekar, A., Collier, A.Y., Chen, B., Barouch, D.H., and Alter, G. (2023). Selective SARS-CoV2 BA.2 escape of antibody Fc/Fc-receptor interactions. iScience 26, 106582.

Beaudoin-Bussieres, G., and Finzi, A. (2024). Deciphering Fc-effector functions against SARS-CoV-2. Trends Microbiol 32, 756–768.

Bok, K., Sitar, S., Graham, B.S., and Mascola, J.R. (2021). Accelerated COVID-19 vaccine development: milestones, lessons, and prospects. Immunity 54, 1636–1651.

Bruhns, P., Iannascoli, B., England, P., Mancardi, D.A., Fernandez, N., Jorieux, S., and Daeron, M. (2009). Specificity and affinity of human Fcgamma receptors and their polymorphic variants for human IgG subclasses. Blood 113, 3716–3725.

Buhre, J.S., Pongracz, T., Kunsting, I., Lixenfeld, A.S., Wang, W., Nouta, J., Lehrian, S., Schmelter, F., Lunding, H.B., Duhring, L., et al. (2022). mRNA vaccines against SARS-CoV-2 induce comparably low long-term IgG Fc galactosylation and sialylation levels but increasing long-term IgG4 responses compared to an adenovirus-based vaccine. Front Immunol 13, 1020844.

Cassel, D.L., Keller, M.A., Surrey, S., Schwartz, E., Schreiber, A.D., Rappaport, E.F., and McKenzie, S.E. (1993). Differential expression of Fc gamma RIIA, Fc gamma RIIB and Fc gamma RIIC in hematopoietic cells: analysis of transcripts. Mol Immunol 30, 451–460.

Chan, C.E.Z., Seah, S.G.K., Chye, H., Massey, S., Torres, M., Lim, A.P.C., Wong, S.K.K., Neo, J.J.Y., Wong, P.S., Lim, J.H., et al. (2021). The Fc-mediated effector functions of a potent SARS-CoV-2 neutralizing antibody, SC31, isolated from an early convalescent COVID-19 patient, are essential for the optimal therapeutic efficacy of the antibody. PLoS One 16, e0253487.

Clark, J.J., Hoxie, I., Adelsberg, D.C., Sapse, I.A., Andreata-Santos, R., Yong, J.S., Amanat, F., Tcheou, J., Raskin, A., Singh, G., et al. (2024). Protective effect and molecular mechanisms of human non-neutralizing cross-reactive spike antibodies elicited by SARS-CoV-2 mRNA vaccination. Cell Rep 43, 114922.

DiLillo, D.J., Tan, G.S., Palese, P., and Ravetch, J.V. (2014). Broadly neutralizing hemagglutinin stalk-specific antibodies require FcgammaR interactions for protection against influenza virus in vivo. Nat Med 20, 143–151.

Fielding, C.A., Sabberwal, P., Williamson, J.C., Greenwood, E.J.D., Crozier, T.W.M., Zelek, W., Seow, J., Graham, C., Huettner, I., Edgeworth, J.D., et al. (2022). SARS-CoV-2 host-shutoff impacts innate NK cell functions, but antibody-dependent NK activity is strongly activated through non-spike antibodies. Elife 11.

Fleit, H.B., and Kobasiuk, C.D. (1991). The human monocyte-like cell line THP-1 expresses Fc gamma RI and Fc gamma RII. J Leukoc Biol 49, 556–565.

Gaebler, C., Wang, Z., Lorenzi, J.C.C., Muecksch, F., Finkin, S., Tokuyama, M., Cho, A., Jankovic, M., Schaefer-Babajew, D., Oliveira, T.Y., et al. (2021). Evolution of antibody immunity to SARS-CoV-2. Nature 591, 639–644.

Gelderloos, A.T., Verheul, M.K., Middelhof, I., de Zeeuw-Brouwer, M.L., van Binnendijk, R.S., Buisman, A.M., and van Kasteren, P.B. (2024). Repeated COVID-19 mRNA vaccination results in IgG4 class switching and decreased NK cell activation by S1-specific antibodies in older adults. Immun Ageing 21, 63.

Goldblatt, D., Alter, G., Crotty, S., and Plotkin, S.A. (2022). Correlates of protection against SARS-CoV-2 infection and COVID-19 disease. Immunol Rev 310, 6–26.

Gorman, M.J., Patel, N., Guebre-Xabier, M., Zhu, A.L., Atyeo, C., Pullen, K.M., Loos, C., Goez-Gazi, Y., Carrion, R., Jr., Tian, J.H., et al. (2021). Fab and Fc contribute to maximal protection against SARS-CoV-2 following NVX-CoV2373 subunit vaccine with Matrix-M vaccination. Cell Rep Med 2, 100405.

Graham, C., Lechmere, T., Rehman, A., Seow, J., Kurshan, A., Huettner, I., Maguire, T.J.A., Tam, J.C.H., Cox, D., Ward, C., et al. (2022). The effect of Omicron breakthrough infection and extended BNT162b2 booster dosing on neutralization breadth against SARS-CoV-2 variants of concern. PLoS Pathog 18, e1010882.

Graham, C., Seow, J., Huettner, I., Khan, H., Kouphou, N., Acors, S., Winstone, H., Pickering, S., Galao, R.P., Dupont, L., et al. (2021). Neutralization potency of monoclonal antibodies recognizing dominant and subdominant epitopes on SARS-CoV-2 Spike is impacted by the B.1.1.7 variant. Immunity 54, 1276–1289.

Hartley, G.E., Fryer, H.A., Gill, P.A., Boo, I., Bornheimer, S.J., Hogarth, P.M., Drummer, H.E., O’Hehir, R.E., Edwards, E.S.J., and van Zelm, M.C. (2024). Homologous but not heterologous COVID-19 vaccine booster elicits IgG4+ B-cells and enhanced Omicron subvariant binding. NPJ Vaccines 9, 129.

Higdon, M.M., Wahl, B., Jones, C.B., Rosen, J.G., Truelove, S.A., Baidya, A., Nande, A.A., ShamaeiZadeh, P.A., Walter, K.K., Feikin, D.R., et al. (2022). A Systematic Review of Coronavirus Disease 2019 Vaccine Efficacy and Effectiveness Against Severe Acute Respiratory Syndrome Coronavirus 2 Infection and Disease. Open Forum Infect Dis 9, ofac138.

Howarth, M., Liu, W., Puthenveetil, S., Zheng, Y., Marshall, L.F., Schmidt, M.M., Wittrup, K.D., Bawendi, M.G., and Ting, A.Y. (2008). Monovalent, reduced-size quantum dots for imaging receptors on living cells. Nat Methods 5, 397–399.

Irrgang, P., Gerling, J., Kocher, K., Lapuente, D., Steininger, P., Habenicht, K., Wytopil, M., Beileke, S., Schäfer, S., Zhong, J., et al. (2023). Class switch toward noninflammatory, spike-specific IgG4 antibodies after repeated SARS-CoV-2 mRNA vaccination. Science Immunology 8.

Jackson, C.B., Farzan, M., Chen, B., and Choe, H. (2022). Mechanisms of SARS-CoV-2 entry into cells. Nat Rev Mol Cell Biol 23, 3–20.

Kim, W., Zhou, J.Q., Horvath, S.C., Schmitz, A.J., Sturtz, A.J., Lei, T., Liu, Z., Kalaidina, E., Thapa, M., Alsoussi, W.B., et al. (2022). Germinal centre-driven maturation of B cell response to mRNA vaccination. Nature 604, 141–145.

Kirsebom, F.C.M., Andrews, N., Stowe, J., Ramsay, M., and Lopez Bernal, J. (2023). Duration of protection of ancestral-strain monovalent vaccines and effectiveness of bivalent BA.1 boosters against COVID-19 hospitalisation in England: a test-negative case-control study. Lancet Infect Dis 23, 1235–1243.

Lee, I.T., Cosgrove, C.A., Moore, P., Bethune, C., Nally, R., Bula, M., Kalra, P.A., Clark, R., Dargan, P.I., Boffito, M., et al. (2023). Omicron BA.1-containing mRNA-1273 boosters compared with the original COVID-19 vaccine in the UK: a randomised, observer-blind, active-controlled trial. Lancet Infect Dis 23, 1007–1019.

Mercado, N.B., Zahn, R., Wegmann, F., Loos, C., Chandrashekar, A., Yu, J., Liu, J., Peter, L., McMahan, K., Tostanoski, L.H., et al. (2020). Single-shot Ad26 vaccine protects against SARS-CoV-2 in rhesus macaques. Nature 586, 583–588.

Motta, R.V., and Culver, E.L. (2024). IgG4 autoantibodies and autoantigens in the context of IgG4-autoimmune disease and IgG4-related disease. Front Immunol 15, 1272084.

Mozdzanowska, K., Feng, J., and Gerhard, W. (2003). Virus-neutralizing activity mediated by the Fab fragment of a hemagglutinin-specific antibody is sufficient for the resolution of influenza virus infection in SCID mice. J Virol 77, 8322–8328.

Muecksch, F., Wang, Z., Cho, A., Gaebler, C., Ben Tanfous, T., DaSilva, J., Bednarski, E., Ramos, V., Zong, S., Johnson, B., et al. (2022). Increased Memory B Cell Potency and Breadth After a SARS-CoV-2 mRNA Boost. Nature.

Olivieri, D.N., and Gambon Deza, F. (2018). Immunoglobulin genes in Primates. Mol Immunol 101, 353–363.

Parhiz, H., Atochina-Vasserman, E.N., and Weissman, D. (2024). mRNA-based therapeutics: looking beyond COVID-19 vaccines. Lancet 403, 1192–1204.

Pierre, C.N., Adams, L.E., Higgins, J.S., Anasti, K., Goodman, D., Mielke, D., Stanfield-Oakley, S., Powers, J.M., Li, D., Rountree, W., et al. (2024). Non-neutralizing SARS-CoV-2 N-terminal domain antibodies protect mice against severe disease using Fc-mediated effector functions. PLoS Pathog 20, e1011569.

Pillai, S. (2023). Is it bad, is it good, or is IgG4 just misunderstood? Sci Immunol 8, eadg7327.

Polack, F.P., Thomas, S.J., Kitchin, N., Absalon, J., Gurtman, A., Lockhart, S., Perez, J.L., Perez Marc, G., Moreira, E.D., Zerbini, C., et al. (2020). Safety and Efficacy of the BNT162b2 mRNA Covid-19 Vaccine. N Engl J Med 383, 2603–2615.

Polycarpou, A.e.a. (2023). The complement pattern recognition molecule CL-11 promotes invasion and injury of respiratory epithelial cells by SARS-CoV-2. bioRxiv.

Qin, L., Tang, L.F., Cheng, L., and Wang, H.Y. (2022). The clinical significance of allergen-specific IgG4 in allergic diseases. Front Immunol 13, 1032909.

Rispens, T., and Huijbers, M.G. (2023). The unique properties of IgG4 and its roles in health and disease. Nat Rev Immunol 23, 763–778.

Roltgen, K., Nielsen, S.C.A., Silva, O., Younes, S.F., Zaslavsky, M., Costales, C., Yang, F., Wirz, O.F., Solis, D., Hoh, R.A., et al. (2022). Immune imprinting, breadth of variant recognition, and germinal center response in human SARS-CoV-2 infection and vaccination. Cell 185, 1025–1040 e1014.

Rosa, A., Pye, V.E., Graham, C., Muir, L., Seow, J., Ng, K.W., Cook, N.J., Rees-Spear, C., Parker, E., Silva Dos Santos, M., et al. (2021). SARS-CoV-2 can recruit a haem metabolite to evade antibody immunity. Sci Adv 7, eabg7607.

Santos, A.F., James, L.K., Bahnson, H.T., Shamji, M.H., Couto-Francisco, N.C., Islam, S., Houghton, S., Clark, A.T., Stephens, A., Turcanu, V., et al. (2015). IgG4 inhibits peanut-induced basophil and mast cell activation in peanut-tolerant children sensitized to peanut major allergens. J Allergy Clin Immunol 135, 1249–1256.

Schafer, A., Muecksch, F., Lorenzi, J.C.C., Leist, S.R., Cipolla, M., Bournazos, S., Schmidt, F., Maison, R.M., Gazumyan, A., Martinez, D.R., et al. (2021). Antibody potency, effector function, and combinations in protection and therapy for SARS-CoV-2 infection in vivo. J Exp Med 218.

Seow, J., Graham, C., Hallett, S.R., Lechmere, T., Maguire, T.J.A., Huettner, I., Cox, D., Khan, H., Pickering, S., Roberts, R., et al. (2022). ChAdOx1 nCoV-19 vaccine elicits monoclonal antibodies with cross-neutralizing activity against SARS-CoV-2 viral variants. Cell Rep, 110757.

Seow, J., Graham, C., Merrick, B., Acors, S., Pickering, S., Steel, K.J.A., Hemmings, O., O’Byrne, A., Kouphou, N., Galao, R.P., et al. (2020). Longitudinal observation and decline of neutralizing antibody responses in the three months following SARS-CoV-2 infection in humans. Nat Microbiol 5, 1598–1607.

Sokal, A., Chappert, P., Barba-Spaeth, G., Roeser, A., Fourati, S., Azzaoui, I., Vandenberghe, A., Fernandez, I., Meola, A., Bouvier-Alias, M., et al. (2021). Maturation and persistence of the anti-SARS-CoV-2 memory B cell response. Cell 184, 1201–1213 e1214.

Tiller, T., Meffre, E., Yurasov, S., Tsuiji, M., Nussenzweig, M.C., and Wardemann, H. (2008). Efficient generation of monoclonal antibodies from single human B cells by single cell RT-PCR and expression vector cloning. J Immunol Methods 329, 112–124.

Tostanoski, L.H., Wegmann, F., Martinot, A.J., Loos, C., McMahan, K., Mercado, N.B., Yu, J., Chan, C.N., Bondoc, S., Starke, C.E., et al. (2020). Ad26 vaccine protects against SARS-CoV-2 severe clinical disease in hamsters. Nat Med 26, 1694–1700.

Turner, J.S., O’Halloran, J.A., Kalaidina, E., Kim, W., Schmitz, A.J., Zhou, J.Q., Lei, T., Thapa, M., Chen, R.E., Case, J.B., et al. (2021). SARS-CoV-2 mRNA vaccines induce persistent human germinal centre responses. Nature 596, 109–113.

UKHSA (2022). SARS-CoV-2 variants of concern and variants under investigation in England, number 34.

Unger, P.P., Makuch, M., Aalbers, M., Derksen, N.I.L., Ten Brinke, A., Aalberse, R.C., Rispens, T., and van Ham, S.M. (2018). Repeated vaccination with tetanus toxoid of plasma donors with pre-existing specific IgE transiently elevates tetanus-specific IgE but does not induce allergic symptoms. Clin Exp Allergy 48, 479–482.

Urban, M., Winkler, T., Landini, M.P., Britt, W., and Mach, M. (1994). Epitope-specific distribution of IgG subclasses against antigenic domains on glycoproteins of human cytomegalovirus. J Infect Dis 169, 83–90.

van de Veen, W., Globinska, A., Jansen, K., Straumann, A., Kubo, T., Verschoor, D., Wirz, O.F., Castro-Giner, F., Tan, G., Ruckert, B., et al. (2020). A novel proangiogenic B cell subset is increased in cancer and chronic inflammation. Sci Adv 6, eaaz3559.

Van den Hoecke, S., Ehrhardt, K., Kolpe, A., El Bakkouri, K., Deng, L., Grootaert, H., Schoonooghe, S., Smet, A., Bentahir, M., Roose, K., et al. (2017). Hierarchical and Redundant Roles of Activating FcgammaRs in Protection against Influenza Disease by M2e-Specific IgG1 and IgG2a Antibodies. J Virol 91.

van der Neut Kolfschoten, M., Schuurman, J., Losen, M., Bleeker, W.K., Martinez-Martinez, P., Vermeulen, E., den Bleker, T.H., Wiegman, L., Vink, T., Aarden, L.A., et al. (2007). Anti-inflammatory activity of human IgG4 antibodies by dynamic Fab arm exchange. Science 317, 1554–1557.

Vidarsson, G., Dekkers, G., and Rispens, T. (2014). IgG subclasses and allotypes: from structure to effector functions. Frontiers in Immunology 5.

Walls, A.C., Park, Y.J., Tortorici, M.A., Wall, A., McGuire, A.T., and Veesler, D. (2020). Structure, Function, and Antigenicity of the SARS-CoV-2 Spike Glycoprotein. Cell 183, 1735.

Wilson, C., Ebling, R., Henig, C., Adler, T., Nicolaevski, R., Barak, M., Cazabon, J., Maisin, D., Lepoutre, T., Gruson, D., et al. (2013). Significant, quantifiable differences exist between IgG subclass standards WHO67/97 and ERM-DA470k and can result in different interpretation of results. Clin Biochem 46, 1751–1755.

Winkler, E.S., Gilchuk, P., Yu, J., Bailey, A.L., Chen, R.E., Chong, Z., Zost, S.J., Jang, H., Huang, Y., Allen, J.D., et al. (2021). Human neutralizing antibodies against SARS-CoV-2 require intact Fc effector functions for optimal therapeutic protection. Cell 184, 1804–1820 e1816.

Wratil, P.R., Stern, M., Priller, A., Willmann, A., Almanzar, G., Vogel, E., Feuerherd, M., Cheng, C.C., Yazici, S., Christa, C., et al. (2022). Three exposures to the spike protein of SARS-CoV-2 by either infection or vaccination elicit superior neutralizing immunity to all variants of concern. Nat Med 28, 496–503.

Yamin, R., Jones, A.T., Hoffmann, H.H., Schafer, A., Kao, K.S., Francis, R.L., Sheahan, T.P., Baric, R.S., Rice, C.M., Ravetch, J.V., et al. (2021). Fc-engineered antibody therapeutics with improved anti-SARS-CoV-2 efficacy. Nature 599, 465–470.

Zelek, W.M., Harris, C.L., and Morgan, B.P. (2018). Extracting the barbs from complement assays: Identification and optimisation of a safe substitute for traditional buffers. Immunobiology 223, 744–749.

Zhang, A., Stacey, H.D., D’Agostino, M.R., Tugg, Y., Marzok, A., and Miller, M.S. (2023). Beyond neutralization: Fc-dependent antibody effector functions in SARS-CoV-2 infection. Nat Rev Immunol 23, 381–396.

Zohar, T., Atyeo, C., Wolf, C.R., Logue, J.K., Shuey, K., Franko, N., Choi, R.Y., Wald, A., Koelle, D.M., Chu, H.Y., et al. (2022). A multifaceted high-throughput assay for probing antigen-specific antibody-mediated primary monocyte phagocytosis and downstream functions. J Immunol Methods 510, 113328.

Zufferey, R., Nagy, D., Mandel, R.J., Naldini, L., and Trono, D. (1997). Multiply attenuated lentiviral vector achieves efficient gene delivery in vivo. Nat Biotechnol 15, 871–875.

