## Supplemental Information for "Spike-specific IgG4 generated post BNT162b2 mRNA vaccination is inhibitory when directly competing with functional IgG subclasses"

Supplementary Table 1

A)

| Group | Short Group | Extended | 6 Month Waning | Prior Infection | Breakthrough Infection | Bivalent Booster |
|---|---|---|---|---|---|---|
| **Number** | 8 | 17 | 9 | 8 | 9 | 8 |
| **Gender** Male : Female | 1:7 | 7:10 | 4:5 | 3:5 | 3:6 | 1:7 |
| **Age** Range (Median) | 29-57 (46) | 23-66 (47) | 23-66 (47) | 25-58 (33) | 24-58 (29) | 25-66 (48) |

B)

| Group | Mild Infection | Severe Infection |
|---|---|---|
| **Number** | 11 | 14 |
| **Gender** Male : Female | 5:6 | 11:3 |
| **Age** Range (Median) | 27-67* (41)* | 25-83 (55) |

**Supplementary Table 1 – (A)** Demographics of the volunteers in each of the vaccination cohort groups. **(B)** Demographics for volunteers in Mild and Severe Infection groups. *= age data only available for 5 volunteers in this group

Supplementary Figure S1

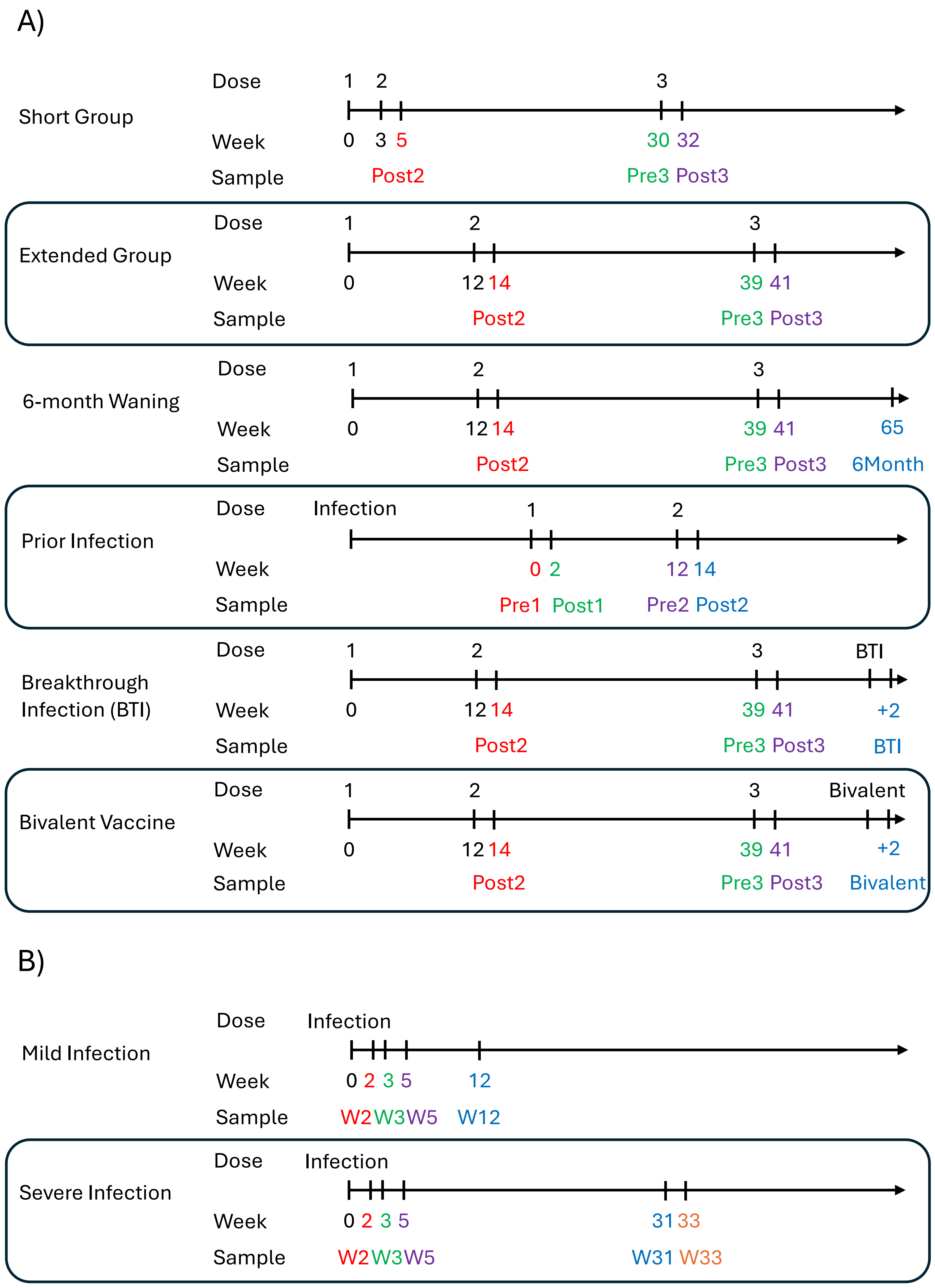

### Supplementary Figure S1

**Supplementary Figure S1 – (A)** Timing of vaccinations and infection in each of our vaccinated groups. Pre1 = before 1$^{st}$ dose, Post1 = 2 weeks after 1$^{st}$ dose, Pre2 = before 2$^{nd}$ dose, Post2 = 2 weeks after 2$^{nd}$ dose, Pre3 = before 3$^{rd}$ dose, Post3 = 2 weeks after 3$^{rd}$ dose, 6Month = 6 months after 3$^{rd}$ dose, BTI = 2 weeks after breakthrough infection after 3$^{rd}$ dose, Bivalent = 2 weeks after bivalent booster. **(B)** Timings of bleed samples from mild and severe infection groups.

### Supplementary Figure S2

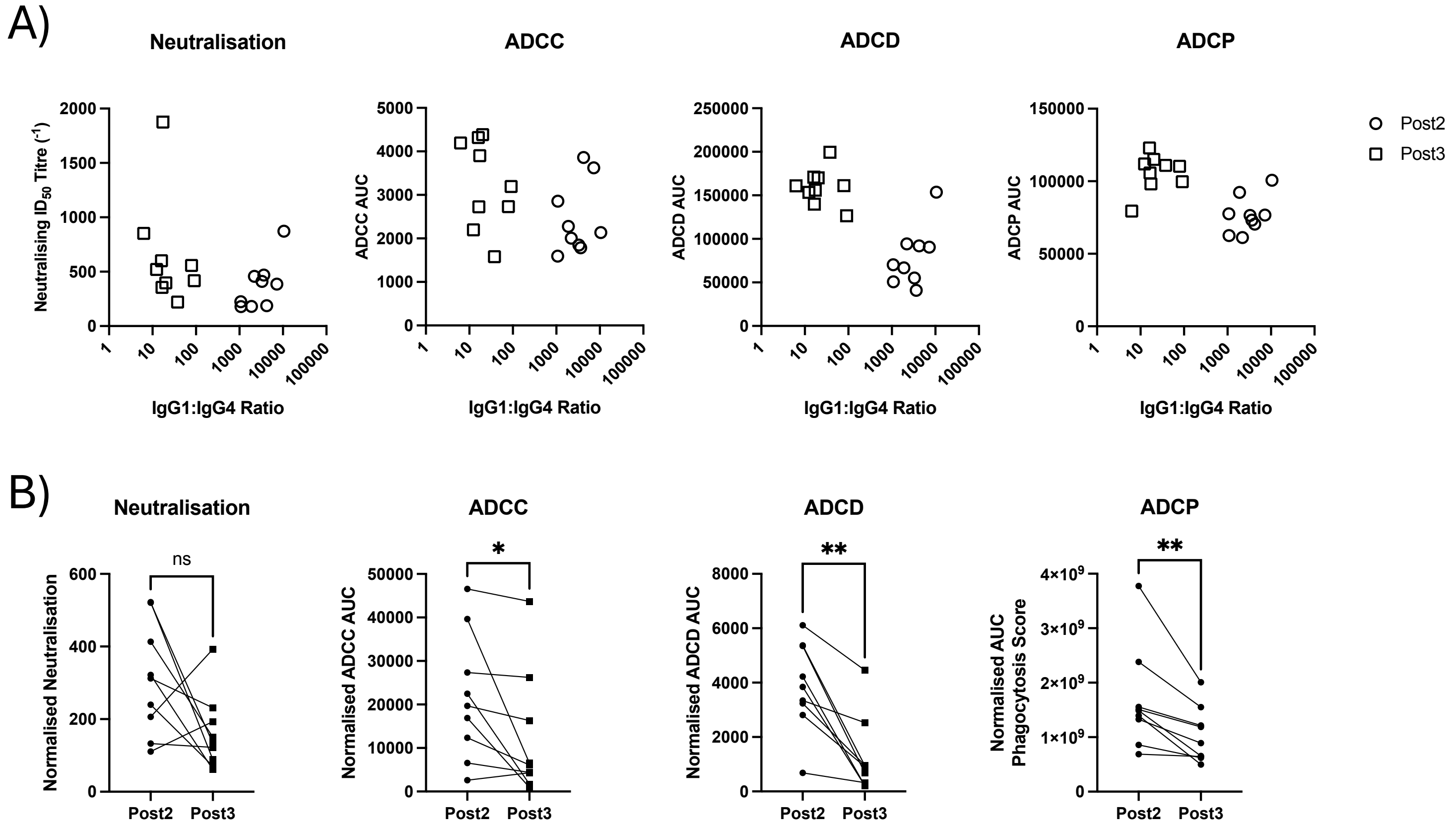


**Supplementary Figure S2 –** (A) Neutralisation, ADCC, ADCD, and ADCP for the extended group, plotted against the IgG1 to IgG4 ratio. Undetectable IgG4 levels were treated as the limit of detection (0.05µg/ml) to calculate values. (B) Normalised Neutralisation, ADCC, ADCD, and ADCP scores calculated by the activity divided by the Total IgG level at either Post2 or Post3 level.
ns not significant, * $p<0.05$, ** $p<0.01$

### Supplementary Figure S3

## A)

| mAb Name | VA14_1 | VA14_R39 | P008_90 | P008_15 | P008_38 | P008_87 | P008_96 | P003_27 | P008_56 | P008_60 | P008_2 | P008_14 | P008_99 | P008_6 |
|---|---|---|---|---|---|---|---|---|---|---|---|---|---|---|
| Specificity | RBD | RBD | RBD | RBD | RBD | RBD | RBD | NTD | NTD | Non-S1 | NTD | NTD | NTD | Non-S1 |
| Competition Group | 1 | 1 | 3 | 4 | 4 | 4 | 4 | 6 | 6 | 7 | ND | ND | ND | ND |
| Antibody Class | 4 | 4 | 1 / 2 | 3 | 3 | 3 | 3 | - | - | - | - | - | - | - |
| WT Spike ELISA EC50 (µg/ml) | 0.0358 | 0.0585 | 0.0182 | 0.0337 | 0.0578 | 0.0212 | 0.0299 | 0.0039 | 0.0602 | 0.0311 | 0.0084 | 0.0662 | 0.0222 | 0.1 |
| WT SARS-CoV-2 Neutralisation IC50 (µg/ml) | 7.34 | 0.07 | 0.0709 | 0.0553 | 0.11 | 4.96 | 9.09 | 1.64 | 0.0136 | 32.9 | >100 | 1.81 | 0.943 | >100 |

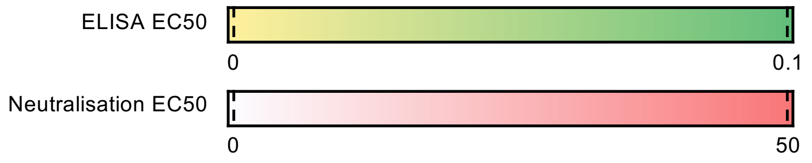


## B)

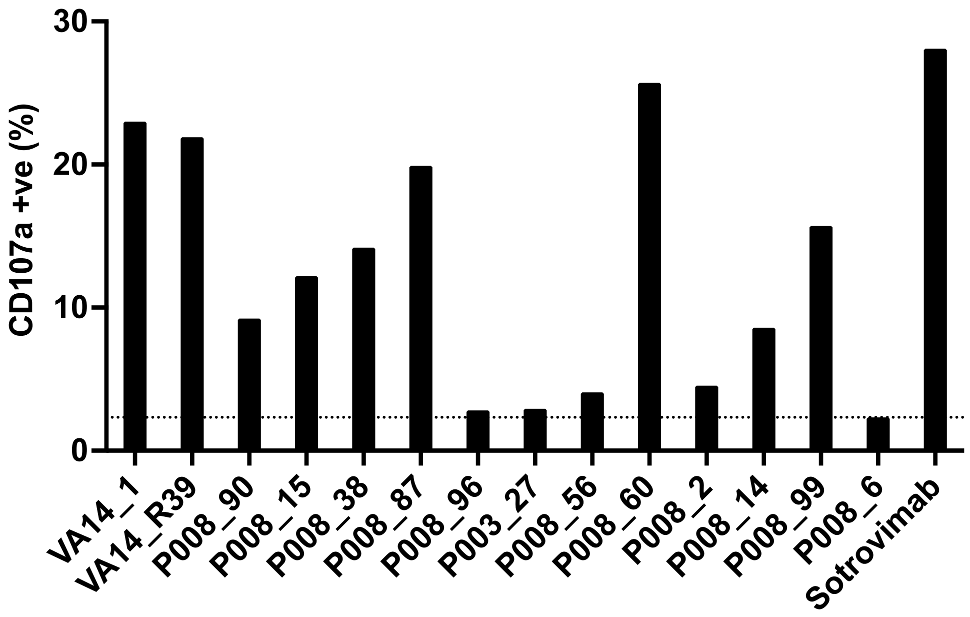


**Supplementary Figure S3 – (A)** Panel of monoclonal antibodies against WT spike, showing their epitope of either Receptor Binding Domain (RBD), N-terminal Domain (NTD), or non-S1 regions; competition group (ND=not done), equivalent antibody class (if an RBD targeting antibody) and the WT spike ELISA EC50 and WT SARS-CoV-2 Pseudovirus Neutralisation IC50. **(B)** ADCC assay measuring activation of NK cells by surface expression of CD107a against same panel of monoclonal antibodies.

Supplementary Figure S4

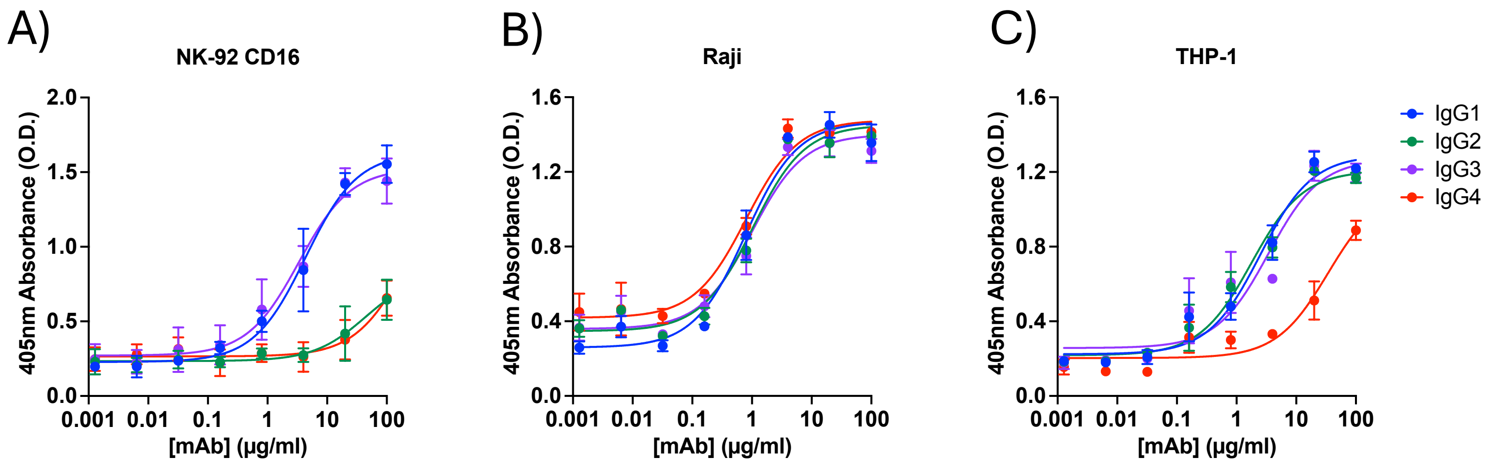


**Supplementary Figure S4 –** Binding of **i**mmune complexes formed pooled monoclonal IgG1 (blue), IgG2 (green), IgG3 (purple), and IgG4 (red) incubated with recombinant WT Spike, on NK cells **(A)**, Raji cells **(B)**, and differentiated THP-1 cells **(C)**. Data is from 3 biological replicates, plotting geometric mean and standard deviation, and representative of 2 technical replicates.

### Supplementary Figure S5

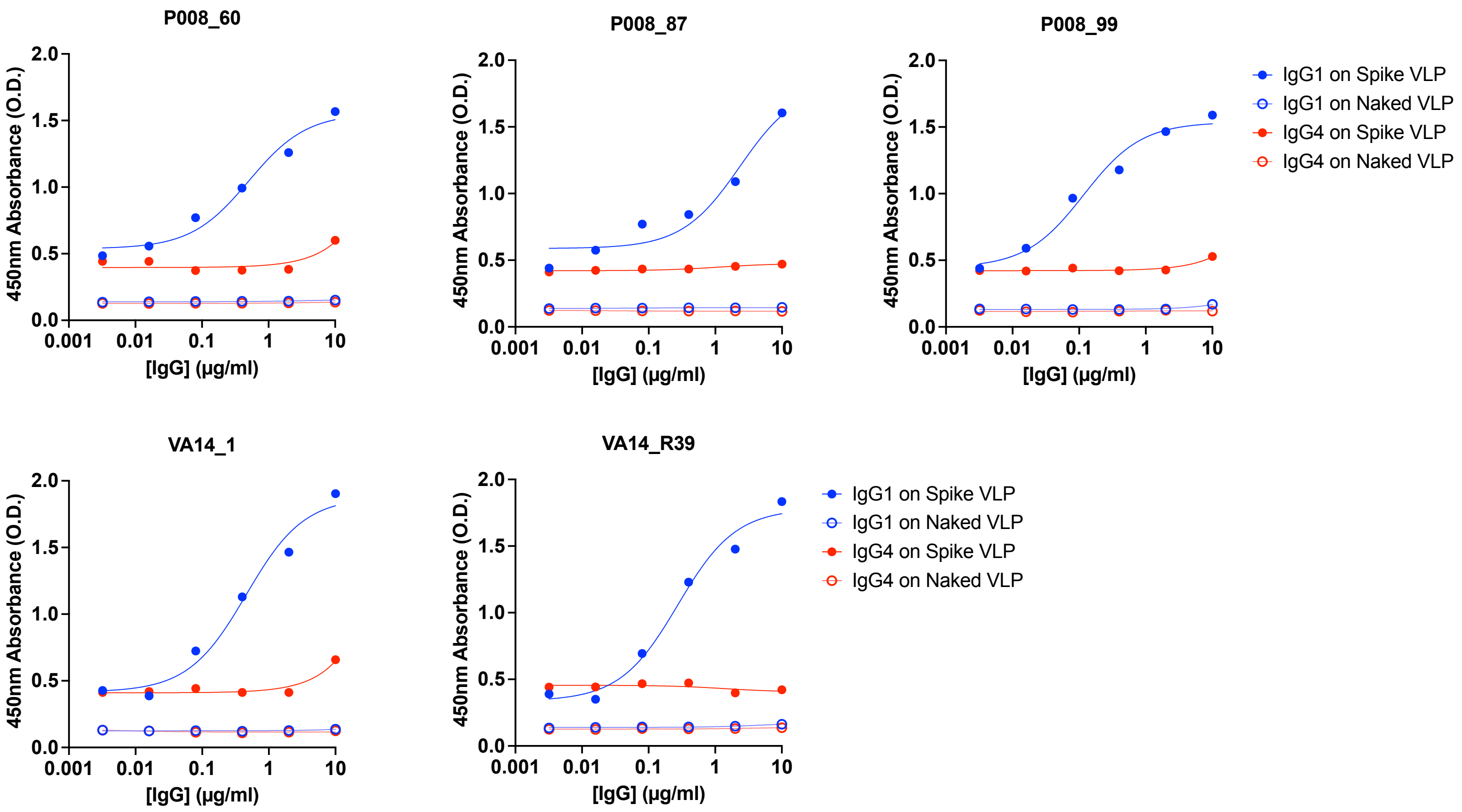


**Supplementary Figure S5** – ADCD assay, measuring Complement C3 deposition stimulated by monoclonal IgG1 (blue) or IgG4 (red), against WT Spike Virus-like Particles (VLP) (filled circles) or Naked VLPs (clear circles).

### Supplementary Figure S6

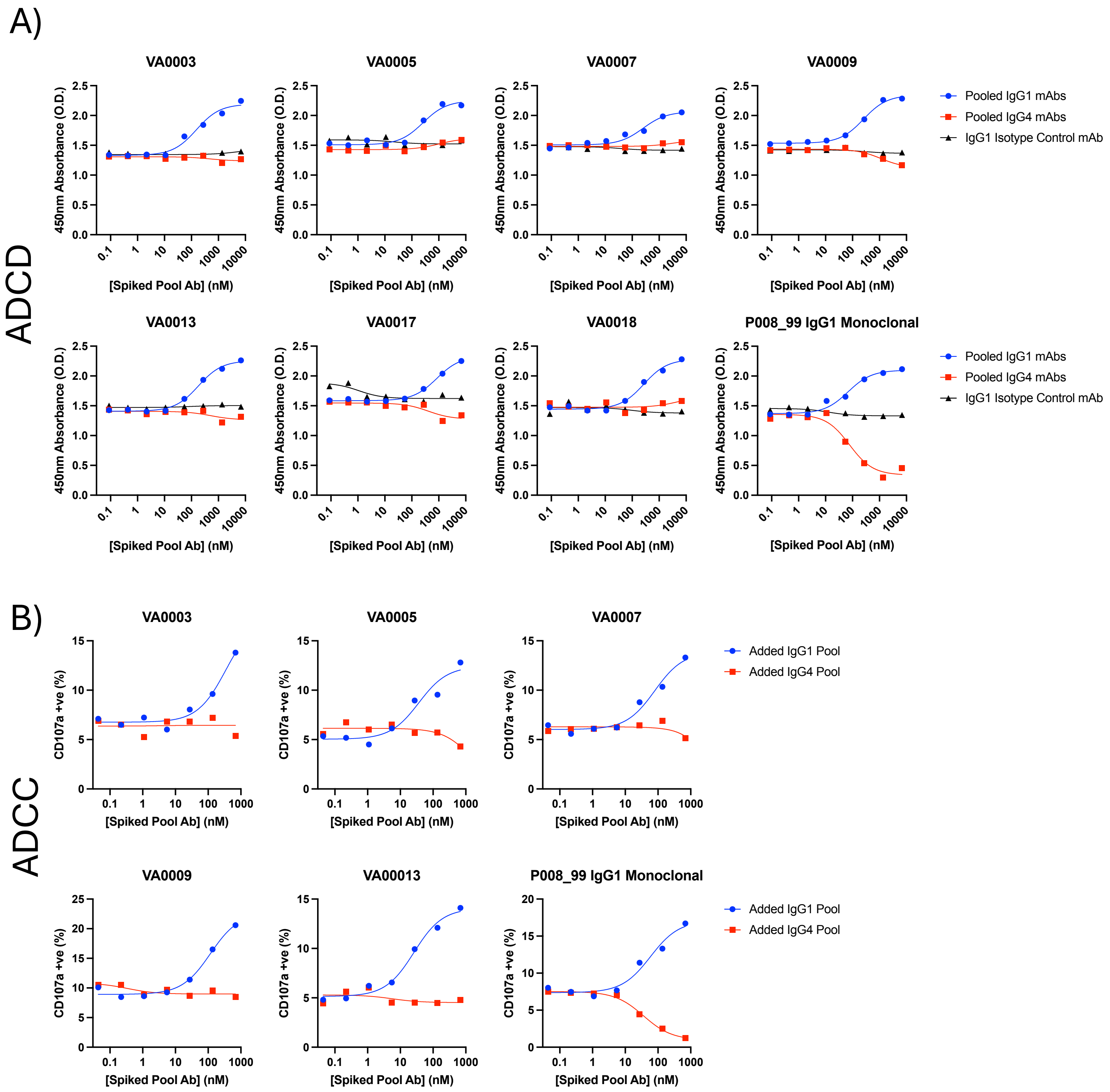


**Supplementary Figure S6 – (A)** ADCD assay spiking in additional Pooled IgG1 or IgG4 monoclonals (P008_60, P008_87, P008_99, VA14_1 and VA14_R39) or IgG1 Isotype Control antibody (anti-HIV IgG1 mAb PGT128) into 7 volunteer plasma samples (Extended Group Post2 samples) at 1:100 dilution or positive control of P008_99 IgG1 monoclonal at 1µg/ml. (**B**) ADCC assay spiking in additional Pooled IgG1 or IgG4 monoclonals into 5 volunteer plasma samples (Extended Group Post2 samples) at 1:100 dilution or positive control of P008_99 IgG1 monoclonal at 1µg/ml.
